# Uncovering the Gene Regulatory Networks Underlying Macrophage Polarization Through Comparative Analysis of Bulk and Single-Cell Data

**DOI:** 10.1101/2021.01.20.427499

**Authors:** Klebea Carvalho, Elisabeth Rebboah, Camden Jansen, Katherine Williams, Andrew Dowey, Cassandra McGill, Ali Mortazavi

## Abstract

Gene regulatory networks (GRNs) provide a powerful framework for studying cellular differentiation. However, it is less clear how GRNs encode cellular responses to everyday microenvironmental cues. Macrophages can be polarized and potentially repolarized based on environmental signaling. In order to identify the GRNs that drive macrophage polarization and the heterogeneous single-cell subpopulations that are present in the process, we used a high-resolution time course of bulk and single-cell RNA-seq and ATAC-seq assays of HL-60-derived macrophages polarized towards M1 or M2 over 24 hours. We identified transient M1 and M2 markers, including the main transcription factors that underlie polarization, and subpopulations of naive, transitional, and terminally polarized macrophages. We built bulk and single-cell polarization GRNs to compare the recovered interactions and found that each technology recovered only a subset of known interactions. Our data provide a resource to study the GRN of cellular maturation in response to microenvironmental stimuli in a variety of contexts in homeostasis and disease.

## Introduction

The developmental programs controlling cellular differentiation are encoded in the genome. The controlled spatiotemporal expression of specific transcription factors (TFs) is at the core of these regulatory events and are hallmarks of cellular differentiation. Each TF interacts transiently with downstream targets that can be described as a dynamic gene regulatory network (GRN) (Peter & Davidson, 2015). The links that make up such networks dictate patterns of gene expression during development and differentiation (Gnanakkumaar et al., 2019; Peter & Davidson, 2011). However, how GRNs apply in the context of cellular maturation in response to microenvironmental stimuli is less clear than in the context of cellular differentiation. While robust GRNs can be built by identifying temporal changes in gene expression and chromatin accessibility using bulk RNA-seq and ATAC-seq data (Duren et al., 2020; Ramirez et al., 2017), whether current single-cell RNA-seq and single-cell ATAC-seq are able to recapitulate or improve bulk-derived GRN connections is unclear. Thus, a side-by-side analysis of GRNs derived using bulk and single-cell techniques in the same model system allows us to quantify the relative strengths of bulk vs single-cell.

GRNs are of particular interest in macrophages, which are able to coordinate immune response to inflammatory conditions, tumors and degenerative disorders (Sharma et al., 2020; Chan & Viswanathan, 2019; DeNardo & Ruffell, 2019). Macrophages are innate immune cells that reside in almost all tissues in the body and play key roles in maintenance of tissue homeostasis and clearance of apoptotic cells (Gordon & Plüddemann, 2018). While heterogeneous, most naive macrophages, termed M0, have the ability to polarize into two main activated states, M1 and M2, based on microenvironmental stimuli (Mills et al., 2000). The ratio of M1:M2 is highly regulated and synchronized in homeostatic tissues (Fujisaka et al., 2009). M1 polarization is induced by bacterial lipopolysaccharides (LPS) and/or the pro-inflammatory cytokine interferon-gamma (IFN-γ) and is therefore generally associated with a pro-inflammatory phenotype, bacterial phagocytosis and anti-tumorigenic activity (Orecchioni et al., 2019; Huang et al., 2018). M2 polarization is induced by interleukins 4 (IL-4) and 13 (IL-13), and is linked to an anti-inflammatory phenotype, helminth resistance and pro-tumorigenic activity (Shaked, 2019; Gordon & Martinez, 2010; Reece et al., 2006). While the M1/M2 framework has been useful to identify major polarization regulatory elements, macrophages *in vivo* may present more complex transcriptional signatures. Multiple M2-like subtypes with different gene expression profiles have been described such as M2a, M2b, M2c and M2d, which are determined by distinct inducing stimuli (Rőszer, 2015). Nonetheless, the M1/M2 dogma of polarization is still a useful conceptual framework to illustrate the main functions as well as the regulatory mechanisms of both major groups (Budhu et al., 2021; Leonard et al., 2020; Bok et al., 2018).

The macrophage phenotype present in a microenvironment can be used as a predictor of disease prognosis. For example, M1 plays an important role in promoting potentially fatal cytokine storms in high risk patients with COVID19 (Lara et al., 2020). Also, prolonged activation of resident macrophages of the brain (microglia) increases both amyloid and tau pathology and may be linked to Alzheimer’s disease (AD) pathogenesis (Kinney et al., 2018). The potential role of activated macrophages in disease has motivated the search for drugs to control polarization (Zhang et al., 2019; Guerriero, 2018). One cancer treatment in development attempts to switch the phenotype of M2-like tumor associated macrophages to an anti-tumorigenic M1 phenotype, highlighting M2’s ability to repolarize towards M1 (Zheng et al., 2017). However, little is known about the transcriptome profile of M2 repolarized towards M1. Although progress has been made in understanding gene expression in terminally polarized M1 and M2 subtypes (Orecchioni et al., 2019), temporal transcriptional changes that drive polarization and repolarization remain poorly understood. Thus, reliable transient polarization markers are needed (Walentynowicz et al., 2018). Recent studies demonstrated that the TFs STAT1 and IRF7 are upregulated in M1, whereas STAT6 is upregulated in M2 both *in vivo* and *in vitro* (Orecchioni et al., 2019; Yu et al., 2019). Albeit relevant, most of the current polarization studies focus mainly on changes in the STAT family of TFs, perhaps overlooking other transcriptional regulators (Ding et al., 2019). Furthermore, no single-cell study to date has clearly captured the heterogeneity of the spectrum of cells that undergo M0 to M1 or M2 activation.

Promoting or inhibiting M0 polarization towards M1 or M2 requires insight into changes to the chromatin landscape and gene expression that precede cell specialization (Briggs et al., 2018; Buenrostro et al., 2018). A useful framework to study macrophage differentiation is the HL-60-derived human macrophage model induced with phorbol-12-myristate-13-acetate (PMA) (Dao et al., 2020; Ramirez et al., 2017; Murao et al., 1983). A considerable amount of gene expression and chromatin accessibility data has been generated using the HL-60 model system (Wenzel et al., 2020; Antwi et al., 2020; Cusanovich et al., 2015; Poplutz et al., 2014). We have previously used bulk RNA-seq and ATAC-seq to build GRNs of HL-60 differentiation into M0s (Ramirez et al., 2017). Thus, profiling mRNA expression and chromatin accessibility during the transition from HL-60-derived-M0 to M1 or M2 with fine time resolution is a valuable approach to identify the genomic mechanisms that drive macrophage polarization (Chistiakov et al., 2018). Last but not least, with the increasing adoption and throughput of single-cell techniques, we can now compare in a well-defined setting the quality of the networks derived using bulk or single-cell techniques.

Here, we characterized the genomic regulatory events promoting macrophage polarization using the macrophage-differentiated HL-60 model to identify cell state transitions, intermediary markers and the gene regulatory networks between the initial M0 pre-stimulus and the terminally polarized M1 and M2 states. Our approach profiled the dynamic changes in the transcriptome and the chromatin landscape at bulk and single-cell levels at 3, 6, 12, and 24 hours of macrophage polarization. Furthermore, we explored the transcriptome profile of M2 repolarized towards M1. We identified key TFs at the core of the regulatory pathways that control polarization into M1 and M2 states and M2 to M1 repolarization. We built GRNs using bulk and single-cell data, and we validated the targets of multiple TFs in M2 polarization. Finally, we compared the GRNs that we derived separately using bulk and single-cell data to identify what portions of the GRNs are recovered by either or both sets of methods.

## Results

### Distinct subsets of genes drive macrophage polarization towards M1 or M2 states

We activated HL-60-derived M0 either to M1 with LPS and IFN-γ or to M2 with IL-13 and IL-4, respectively, in order to identify how subtype-specific polarization affects macrophage gene expression. We collected samples for bulk RNA-seq at 3, 6, 12, and 24 hours post-stimulation (Figures 1A and S1A). We identified 7,601 genes (alpha < 0.05, FDR < 0.05%) whose expressions vary in a time-specific fashion using maSigPro. These genes grouped into 18 distinct clusters from which we selected 11 clusters (4,760 genes) representing four major patterns of expression for HL-60-, M0-, M1-, and M2-specific responses (Figure 1B). Each cluster contains distinct subtype-specific signaling molecules and TFs (Figure 1B). We identified 1,269 genes whose expression is higher in M1 (clusters Rc2, Rc5, Rc6 and Rc15) and 1,462 genes whose expression is higher in M2 (clusters Rc1, Rc7 and Rc13). We used an UpSet plot of maSigPro detected genes to identify a subset of 1,194 genes that had a similarly increased expression in both M1 and M2, and another subset of 149 genes that had a higher expression in HL-60 and M2 only (Figure 1C). These subtype specific genes showed distinct temporal expression patterns (Figures 1D & S1B). Canonical M1 polarization markers such as *CXCL10, CXCL11* and *GBP4* (Tang et al., 2017; Mantovani et al., 2006) increased expression rapidly by 3 hours post-stimuli and peaked expression around 6 hours (Figure 1D Upper). Canonical M2-associated genes *CCL24* and *CLEC4A* (Makita et al., 2015; Tang et al., 2017) were also induced at 3 hours post-stimuli and reached peak expression around 6 hours (Figure 1D Lower). M2 also showed higher expression of signaling molecule *CSF2* that was recently shown to promote macrophages transition into an M2 phenotype (Li et al., 2020). Therefore, important expression changes leading to terminal polarization of macrophage subtypes were established within the first 6 hours following addition of stimuli. We also identified a subset of pro-inflammatory chemokines present in M1-specific clusters, such as *CCL7, CCL8, CXCL9, CXCL10*, and *CXCL11* (Gurvich et al., 2020; Lu et al., 2018) (Figure S1C). M2 clusters showed higher expression of chemokines *CCL24*, which is upregulated in macrophages stimulated with IL-4 (Lee et al., 2020) and *CKLF*, which has been associated with decreasing inflammation in dermal disorders (Zheng et al., 2017). Overall, we found substantial and rapid differences in gene expression as the result of macrophage polarization.

**Figure 1.**
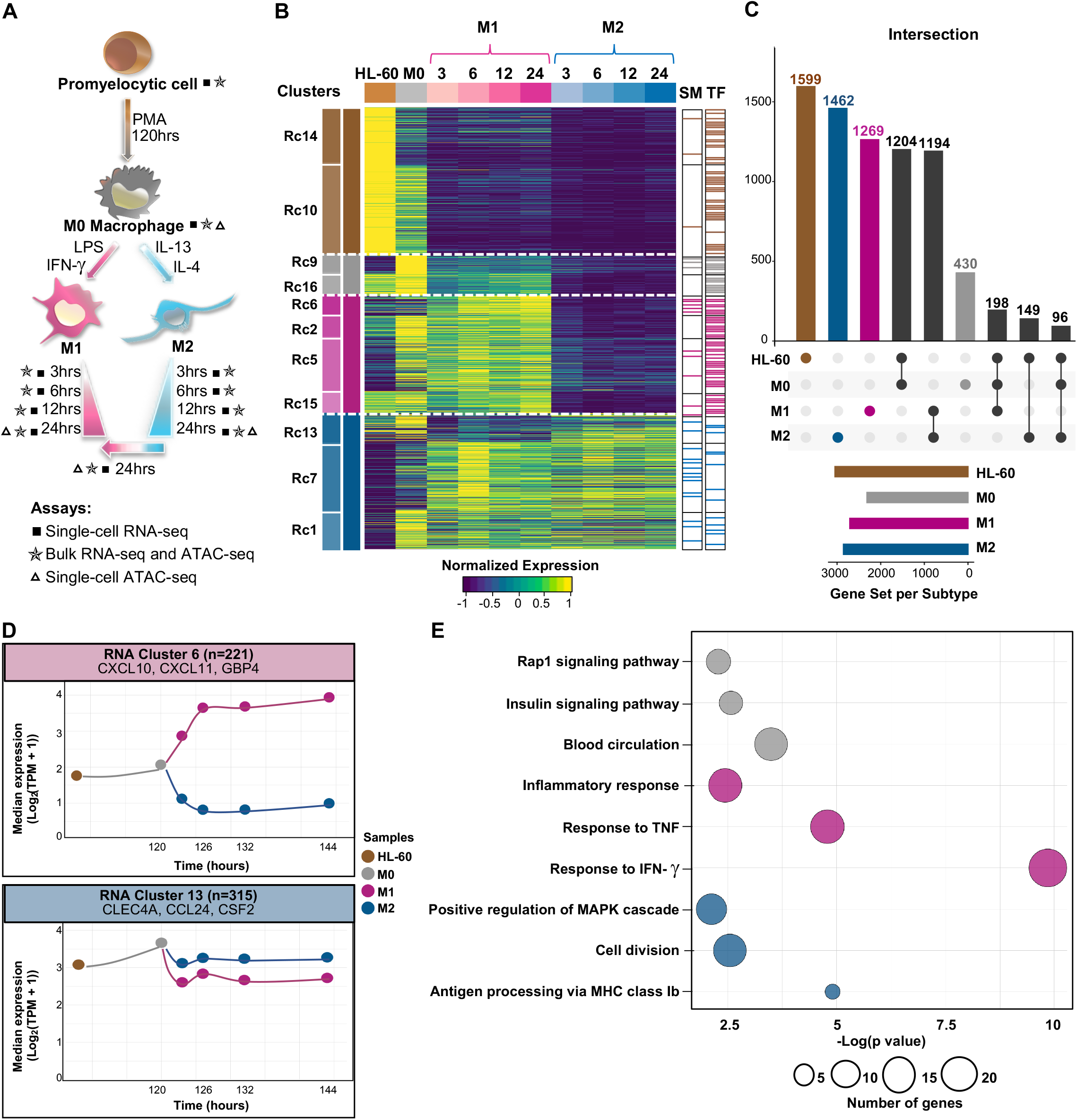
HL-60-derived M0 polarization reveals distinct clusters of M1- and M2-specific genes. A) Schematic diagram of experimental design highlighting samples processed and distinct assays performed. B) Heatmap of 4,760 genes with dynamic temporal profiles identified by MaSigPro clustering (alpha < 0.05, FDR < 0.05%). Each column represents the average expression for a time point and each row represents a gene. Each cluster represents a subset of genes that show a similar pattern of expression along the time course. Brown, grey, pink, and blue represent HL-60-, M0-, M1-, and M2-specific clusters, respectively. RNA-seq data (TPM) is row-mean normalized. Signaling molecules (SM) and transcription factors (TF) present in each cluster are shown.C) UpSet plot highlights distinct and overlapping genes across main subtypes (HL-60, M0, M1, and M2). D) Representative cluster of 221 genes (RNA Cluster 6) that exhibit increased expression during the M1 polarization time course. Representative cluster of 315 gen E) Gene ontology (GO) enrichment analyses of M0 (430), M1 (1269), and M2 (1462) differentially regulated genes.

We performed gene ontology analysis on M0-, M1-, and M2-specific clusters to identify biological processes and pathways associated with subtype specific genes (Figure 1E). M0-specific genes are enriched for Rap1 and insulin signaling pathways, which are important for macrophage response to pathogens and phagocytosis, respectively (Chung et al., 2008; Liang et al., 2007; Rosa et al., 1996).

M1-specific genes are enriched for regulation of inflammatory signaling, such as tumor necrosis factor (TNF) and IFN-γ response, which is a hallmark of M1 macrophages (Figure 1E). Activation of mitogen-activated protein kinase (MAPK) pathway, which is important for cellular proliferation and M2 polarization (Neamatallah, 2019), is enriched in M2-specific clusters. Thus, M2-specific genes are enriched for cellular division (Figure 1E). Interestingly, M2 cells are able to proliferate in our system, whereas M1 are not.

We collected a time course of M1 and M2 polarizing macrophages to identify dynamic temporal expression changes and intermediary markers regulated in response to polarization signaling. 500 genes showed increased expression as early as 3 hours post-stimuli; 400 of those were higher in M1 and 100 higher in M2 (Figure S1D). *FCGR1A*, whose expression is increased in immuno-inflammatory syndromes (Minar et al., 2014), reacted to M1 but not M2 stimuli and was activated as early as 3 hours of polarization. The signaling molecule *CSF2* displayed increased expression at 3 hours during M2 polarization but not during that of M1 (Figure S1D). *CSF2* over-expression leads to increased autophagy that promotes M0 polarization towards an M2 subtype (Chen et al., 2014; Liu et al., 2015). Other known M1-specific genes such as chemokine *CXCL9* and TF GATA2 (Yin et al., 2020) displayed differential expression in intermediary polarization states at 6 and 12 hours post-stimuli, respectively. Intermediary M2 polarization states showed differential expression of *COX6A1* (Codoni et al., 2016), which participates in macrophage oxidative phosphorylation, and M2 TF RXRA (Czimmerer et al., 2018) at 6 and 12 hours of polarization, respectively. Some genes displayed delayed response to stimuli, highlighting the dynamic nature of polarization. For instance, cytokine *CCL7* only showed higher expression in M1 compared to M2 at 24 hours after stimuli, whereas *CLECL1* gene that has been shown to stimulate IL-4 production in T helper cells and is overexpressed in nonclassical monocytes (M2) (Talker et al., 2020) comes up later in M2 polarization at 24 hours. Therefore, distinct subsets of M1- and M2-specific genes were regulated at early, intermediary and late polarization stages.

### Dynamic shifts in chromatin accessibility occur in response to polarization stimuli and are sustained across differentiation

We clustered regions with similar accessibility profiles over time to identify chromatin regions that change during macrophage polarization (Figure 2A). We associated each chromatin region with its nearest gene and identified 25,331 differentially accessible regions (alpha < 0.05, FDR < 0.05%) using MaSigPro. These regions grouped into 21 distinct clusters from which we selected 12 clusters (14,174 regions) representing four major patterns of chromatin accessibility for HL-60-, M0-, M1-, and M2-specifific responses. Eight of these clusters contained regions overall more open in M1 (clusters Ac19, Ac13, Ac11 and Ac5) or M2 (clusters Ac18, Ac6, Ac7 and Ac17) (Figure 2A). The M1-specific cluster Ac13 highlighted regions around genes known to be upregulated in M1, such as apolipoproteins *APOL1, APOL4 and APOL6* (Gurvich et al., 2020; Lee et al., 2020; Mantovani et al., 2006) that became more accessible as M1 polarized and slightly less accessible as M2 polarized (Figure 2B Upper). Cluster Ac18 highlighted regions near known M2-specific genes, such as *STAT6, ADK* and *CCL1* (Mantovani et al., 2006, 2004) that in turn became more accessible during M2 polarization compared to M1 (Figure 2B Lower). Principal component analysis (PCA) revealed that PC2, which accounted for 18.1% of the variance, separated M1 from M2, with M0 sitting at 0 (Figure S2A). Pearson correlation analysis also showed great differences between macrophage subtypes (Figure S2B). Therefore, chromatin landscape reorganization was subtype-specific.

**Figure 2.**
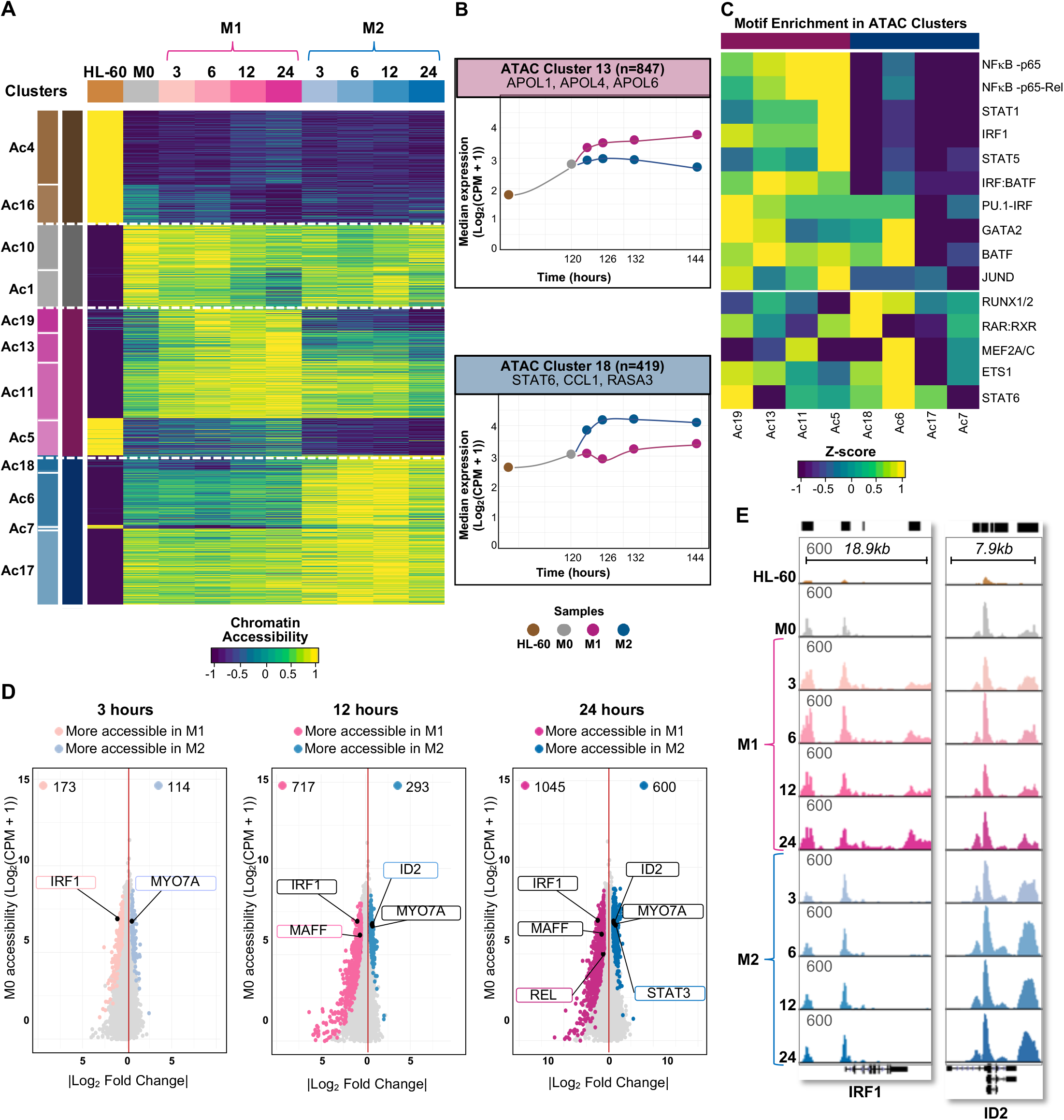
Differential chromatin accessibility occurs early in response to polarization stimuli. A) Heatmap of 14,174 differentially accessible chromatin regions identified by MaSigPro clustering (alpha < 0.05, FDR < 0.05%). Each column represents the average chromatin accessibility for a time point and each row represents a genomic region. Brown, grey, pink, and blue represent HL-60-, M0-, M1-, and M2-specific clusters, respectively. Read counts were TMM normalized, scaled by library size, and row-mean normalized. B) Representative cluster of 847 genomic regions (ATAC Cluster 13) that exhibit increased accessibility during the M1 polarization time course. Representative cluster of 419 genomic regions (ATAC Cluster 18) that exhibit increased accessibility during the M2 polarization time course. C) Hierarchical clustering of known motif enrichment in regions included in the 8 M1- or M2-accessible clusters. D) Genomic regions with differential chromatin accessibility between M1 and M2 (log_2_FC>1, FDR<0.05) compared to normalized M0 accessibility. Genomic regions associated with IRF1, MYO7A, MAFF, ID2, REL, and STAT3 are indicated. E) UCSC genome browser screenshots depicting accessibility around IRF1 and ID2 genes in M1 and M2 samples (y axis scaled from 0 to 600).

We looked for enrichment of TF binding motifs in the M1- or M2-accessible regions of each cluster in order to determine whether a subset of transcription factors drive M1- or M2-specific chromatin changes (Figure 2C). Pro-inflammatory TFs NF_k_B, STAT1, STAT5, and IRF1, which are known to control M1 polarization (Chauhan et al., 2018; Platanitis & Decker, 2018), showed binding motifs enriched in M1 clusters (Figure 2C). JUND, an early target of LPS activation (Srivastava et al., 2013), also showed binding motifs enriched in M1 clusters. In M2 clusters, RAR, STAT6, RUNX and MEF2 motifs were enriched (Figure 2C). While RAR and STAT6 are known to be essential TFs in M2 macrophage activation (Lee et al., 2016), the RUNX and the MEF2 families of TFs are less well-described. RUNX2 has been implicated in the promotion of osteogenic events in both M1 and M2 (Dube et al., 2017; Li et al., 2019) and MEF2 was shown to be repressed by IFN-γ signaling (Kang et al., 2017). Notably, the binding motifs for the RUNX family of TFs are nearly identical, as well as the MEF2 family of TFs have very similar binding motifs. Based on expression, RUNX1 or RUNX2 as well as MEF2A and MEF2C were predicted to bind their family motif match in our data. After scanning for motifs in all open regions, we increased the specificity of our analyses by only scanning for footprints of 8-31nt in regions likely bound by TFs using HINT-ATAC (Li et al., 2019) by pooling the ATAC-seq data for all time points (3 to 24 hours) for M1 (∼350 million reads) or M2 (∼260 million reads). We recovered 362,198 footprints for M1 and 362,809 footprints for M2. The output mostly recapitulated the results from scanning for motifs in clusters of regions identified as M1- or M2-specific seen in Figure 2C. Again, the NF_k_B, JUND, STAT1, and STAT5 TFs had more footprints in M1 than M2; RUNX and MEF2 families of TFs had more footprints in M2 than M1 (Figure S2C). The PU.1-IRF (CGGAAGTGAAAC) and the IRF1 (GAAAGTGAAAGT) motifs were enriched in the M1 subtype (Figure S2C). PU.1-IRF play important roles in macrophage transcriptional response to IFN-γ (Langlais et al., 2016). In summary, we found that distinct subsets of TF are associated with M1 and M2 polarization in our model.

We compared chromatin accessibility between M1 and M2 cells at 3, 12, and 24 hours to investigate temporal changes in the chromatin landscape leading to altered gene regulation upon polarization to M1 or M2 macrophages (Figure 2D). At the 3-hour time point, we detected 173 genes more open in M1 than M2, and 114 genes more open in M2 than M1 (log_2_FC > 1, FDR < 0.05), indicating some early changes in gene accessibility in response to polarization stimuli (Figure 2D). While gene expression changes leading to polarization were established rapidly within the first 6 hours, important chromatin changes occurred later in the time course, after 12 hours (Figures S1D & 2D). M1 stimuli led to increased *IRF1* accessibility (as early as 3 hours of polarization) that was sustained along the time course. IRF1 is known to induce interferon transcription to drive M1 polarization (Platanitis & Decker, 2018). The core macrophage signature gene *MYO7A* (Puranik et al., 2018) became more accessible in M2 at 3 hours and maintained higher accessibility throughout M2 polarization (Figure 2D). TFs MAFF and REL, both induced by LPS signaling (Baillie et al., 2017), became more accessible in M1 than M2 later in the polarization process, at 12 and 24 hours, respectively (Figure 2D). TFs ID2 and STAT3 became more accessible in M2 than M1 at 12 and 24 hours post-stimuli, respectively. *ID2* expression increased greatly and its genomic region became more accessible during M2 polarization (Figures 2D & 2E). Thus, *ID2* is a possible new M2 marker identified in our model.

### Integration of transcriptome and chromatin accessibility dynamics during polarization

Since the bulk ATAC-seq and bulk RNA-seq libraries were prepared from the same pool of cells, their transcriptome profiles and chromatin accessibility landscapes are directly comparable. We hypothesized that tightly regulated genes, such as targets of crucial TFs that share similar expression dynamics, would also share similar accessibility dynamics. In order to compare our previously identified open chromatin and gene expression clusters, we used a Pearson’s χ2 test to determine enrichment between each RNA-seq and ATAC-seq cluster (Figure 3A). Several linked RNA-seq and ATAC-seq clusters show similar dynamics in expression and accessibility. For instance, Rc2 and Ac13 as well as Rc5 and Ac19 had genes with higher expression and associated regions with chromatin more accessible in M1 (Figure 3A). These clusters contained *PU*.*1* and *IRF9*, respectively, which are consistently higher in M1 over M2. M1-enriched clusters also contained *IRF1, CCL8* and *STAT1*. M2-enriched clusters contained *CSF2, MEF2A* and *IL10RA*, which regulate anti-inflammatory function in macrophages (Shouval et al., 2014). Rc1 and Ac6 as well as Rc1 and Ac7 have higher expression and chromatin accessibility in M2 (Figure 3A). These clusters were enriched in *ID2* and M2-specific gene *TIMP1* that is known to respond to M2 cytokine IL-4 (Wang & Joyce, 2010). We therefore detected coordinated changes in chromatin and gene expression for key regulatory factors and polarization markers.

**Figure 3.**
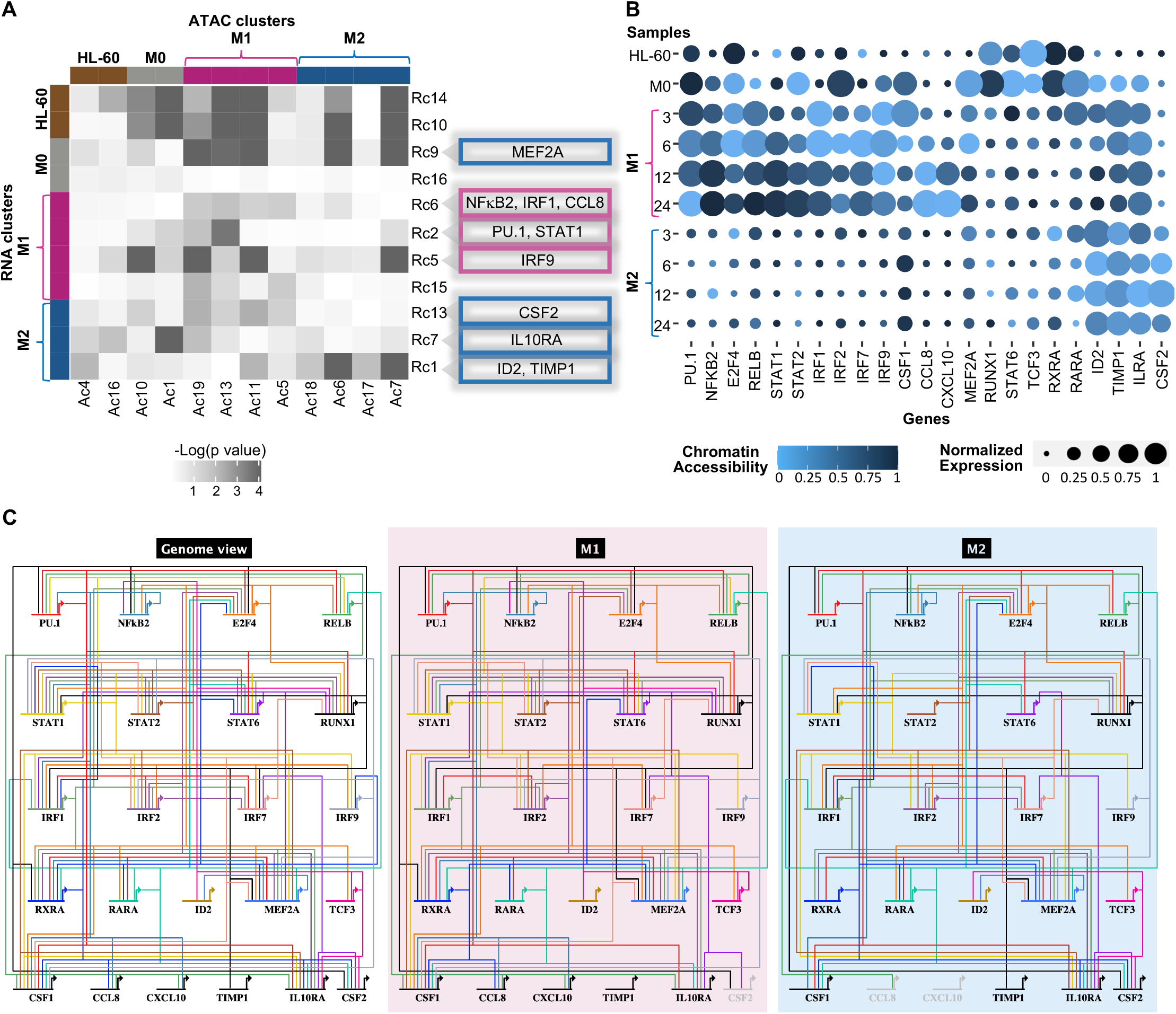
Bulk RNA-seq and ATAC-seq derived gene regulatory networks reveal subtype-specific transcription factor interactions. A)Heatmap of enrichment between each RNA-seq and ATAC-seq cluster calculated using Pearson’s χ2 test. Examples of matching M1-enriched and M2-enriched clusters are indicated in grey. B) Bubble plot of gene expression and chromatin accessibility within the promoter region of key TFs and signaling molecules. Increasing size of the bubble indicates a higher signal in gene expression (TPM scaled across all samples) and a darker blue color indicates higher signal in chromatin accessibility (TMM normalized counts scaled across all samples). C)Gene regulatory networks containing 17 transcription factors and 6 signaling molecules generated from ATAC-seq footprinting and gene expression in bulk RNA-seq data. Genome view shows all connections identified. M1 and M2 panels show the connections specific to each cell type. Each color is specific to a transcription factor that regulates multiple targets.

### Construction of gene regulatory networks from bulk expression and chromatin footprinting data reveals subtype-specific transcription factor interactions

Based on the results above, we narrowed our GRN construction to a set of 17 transcription factors and 6 signaling molecules that stood out in our analyses. *PU*.*1/SPI1, NF*_*k*_*B2, STAT1, STAT2, IRF1, IRF7*, and *IRF9* were highly expressed and more accessible in M1, while *IL10RA, MEF2A, ID2, CSF2*, and *TIMP1* were highly expressed and more accessible in M2 (Figures 3A & 3B). The specific role of PU.1 during macrophage polarization is unclear. However, one study has shown that a microRNA suppressing *PU*.*1* promotes M2 polarization (Li et al., 2018). Another study has demonstrated that knockout of *PU*.*1* leads to a decrease in NF_k_B activation and subsequent inflammation, a M1-specific process (Karpurapu et al., 2011) suggesting that PU.1 plays an important role during M1 polarization.

We merged all time points for M1 and M2 ATAC-seq datasets to achieve >200 million reads needed for chromatin footprinting analysis, as aforementioned. We identified 145,332 footprints in M1 and 148,348 footprints in M2 using the Wellington algorithm (Piper et al., 2013). These footprintings were used to build subtype-specific GRNs (see Methods) focused on our 23 genes of interest represented as circuit diagrams (Figure 3C). At the “genome view” level we identified 141 interactions, which includes 48 total detected in M0 (6 specific), 110 total detected in M1 (40 specific) and 88 total detected in M2 (17 specific); 78 of the 141 connections are shared between some combination of M0/M1/M2. We have previously described the M0 GRN (Ramirez et al., 2017), and therefore focused on M1 and M2 interactions (Figure 3C “M1” & “M2”). As expected, we captured several well-known interactions, such as the PU.1 auto-regulatory feedback loop, RUNX1 regulation of *PU*.*1*, and STAT1 regulation of *IRF1* (Zenke et al., 2018; Lie-A-Ling et al., 2014; Laslo et al., 2006) (Table S1). We decided to also incorporate ID2, which is an inhibitor of some basic helix-loop-helix-containing transcription factors such as *TCF3* (*E2A*) and *TCF12* (*HEB*) even though it does not bind directly to DNA and thus does not have a motif (Rautela et al., 2019). Interestingly, MEF2A binds to *ID2* during M2 polarization, but not during M1 (Figure 3C). Three out of the five TFs targeting *PU*.*1* in our model (PU.1, RUNX1, and RELB) have been confirmed in previous studies (Table S1) and their links to *PU*.*1* are detected in both M1 and M2 GRNs. The targeting of *PU*.*1* by NF_k_B2 was M1-specific. The binding motifs for NF_k_B1 and NF_k_B2 are nearly identical and we selected *NF*_*k*_*B2* in our networks because it is expressed ∼15-fold higher than *NFkB1* in M1. Moreover, *NFkB2* expression is ∼18-fold higher in M1 than in M2 and is therefore an M1-specific TF. This indicates that in our system, the *NFkB* pathway is upregulated specifically in the M1 subtype and plays a key role in regulating other transcription factors such as PU.1. In addition, the binding partner of *NFkB2* (p100), *RELB*, is expressed ∼1.5-fold higher than *RELA* in M1 and *RELB* expression is ∼10-fold higher in M1 than M2. This indicates that in our system, the noncanonical *NFkB* pathway is upregulated specifically in the M1 subtype and plays a key role in regulating other transcription factors such as PU.1 (Figure 3C).

### Single-cell heterogeneity during macrophage polarization reveals distinct activation trajectories

We isolated and sequenced single cells representing each polarization time point seen in Figure 1A using the microfluidic Bio-Rad ddSEQ platform (see Methods) to examine how individual macrophages respond to stimuli. After filtering out low-quality cells, our final dataset contained 18,363 single cells representing subtypes of HL-60 and M0 as well as cells from M1 and M2 polarization time courses (Table S2). A UMAP of 9,993 cells that passed the threshold for the RNA velocity analysis described below showed clear subtype-specific clustering with six distinct cell populations or “Paths” (Figure 4A). HL-60 and M0 cells were enriched in Paths 1 and 2, respectively. Path 3 contained subsets of M1 and M2 time points, whereas Path 4 mainly contained subsets of M0, M1, and M2 cells (Figure 4B). Paths 5 and 6 were enriched for terminally polarized M1 and M2, respectively. Next, we used Monocle to identify a pseudotime course of macrophage polarization (Qiu et al., 2017) to further verify our clustering and reconstruct the main trajectories of these populations (Figure 4C). Paths 5 and 6 were located at the end of two distinct trajectories, thus constituting terminally polarized M1 and M2, respectively. The terminally polarized M1 population contained subsets of all M1 time points (3-, 6-, 12- and 24-hours post-stimuli). Similarly, the terminally polarized M2 population contained subsets of all M2 time points. We thus found that subsets of macrophages respond rapidly to stimuli and our time points contain mixtures of cells at different degrees of polarization.

**Figure 4.**
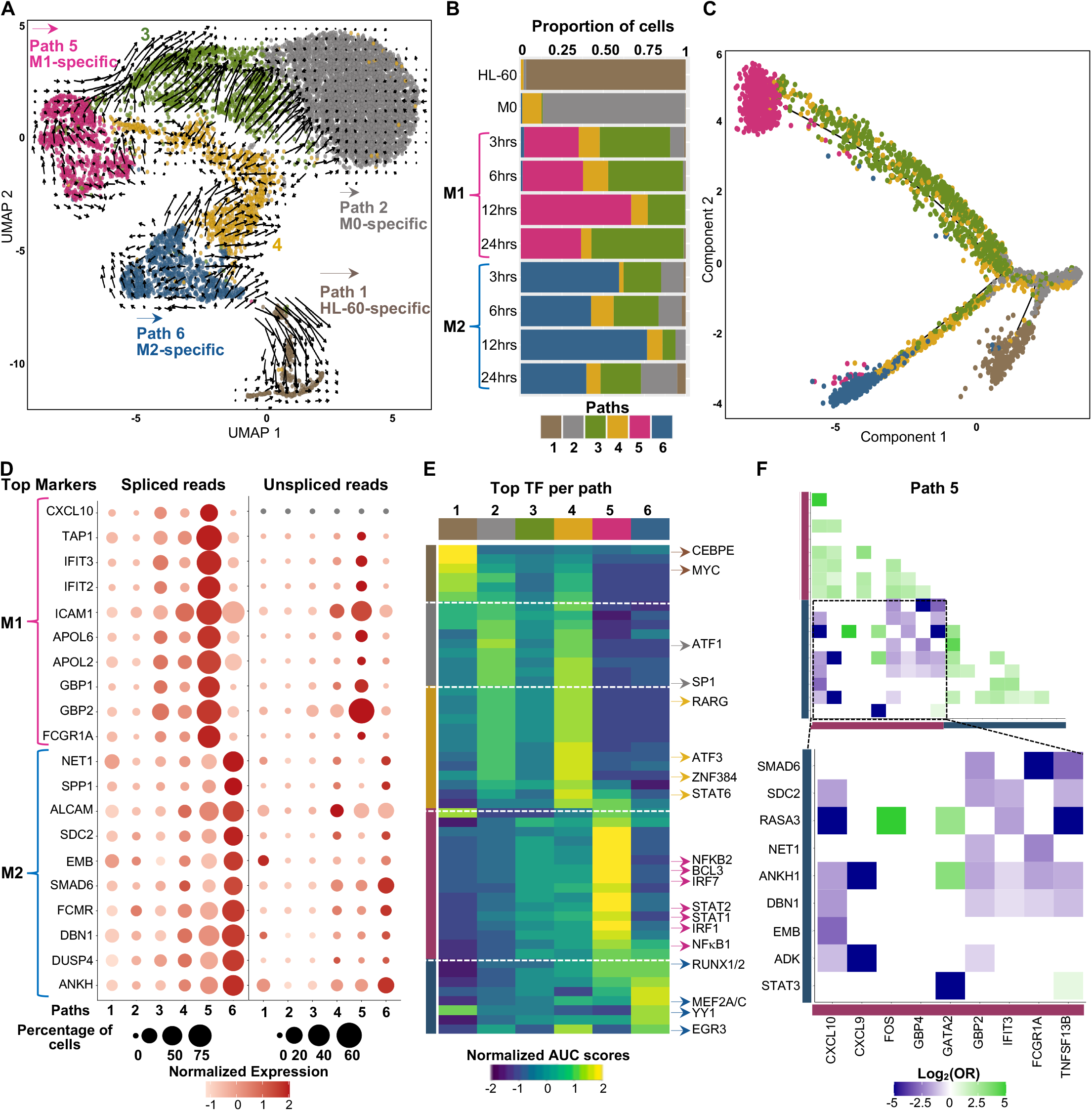
Single-cell RNA-seq and pseudotime analysis identify heterogeneous subpopulations of polarizing macrophages. A)UMAP embedding representation of single-cell RNA-seq polarization time course annotated by clusters of subpopulations (Paths). RNA velocity vectors were projected onto the UMAP and indicate future cellular trajectories. B)Bar plot of the relative proportion of cell subtypes per cluster (Path). C) Pseudotime trajectory of macrophage polarization colored by clusters. Colors are as noted in B. D) Bubble chart depicting spliced and unspliced reads for a given gene (row) per Path (column). Circle sizes represent the percentage of cells expressing the gene. Color key represents normalized average expression. E) Heatmap of AUC scores of transcription factors’ activity per cluster estimated by SCENIC. Columns represent clusters and rows represent top transcription factors that are active per cluster. F) Pairwise odds-ratios (OR) for M1 genes (pink) and M2 genes (blue) detected in Path 5. Odds-ratios that present a p value > 0.05 (calculated by Fisher’s exact test) are set to 0. Data shown as log_2_(odds-ratio).

Paths 3 and 4 were located between M0-M1 and M0-M2 trajectories, suggesting that these cells are transitional cell types (Figure 4C). In addition, we applied RNA velocity (see Methods, Figure S3A) which uses the ratio of unspliced to spliced reads to infer differentiation trajectories represented as a vector whose amplitude and direction depict the future transcriptional state of each cell or group of cells (Figure 4A). This analysis revealed that Path 3 displayed long velocity vectors pointing back towards Path 2 (M0-specific). Moreover, Path 3’s unspliced signal for M1 and M2 markers were similar to Path 2’s (M0) unspliced signal for those markers, indicating that Path 3 was regressing towards the naive M0 state (Figure 4D). This suggests that Path 3 represented macrophages that do not respond to stimuli and tend to maintain a naive state. We observed that a subset of Path 4 cells was progressing towards Path 5 (M1-specific) and another subset was moving towards Path 6 (M2-specific) based on arrow directions and cluster proximity. Path 4 also showed unspliced reads signal of both M1 and M2 markers (Figure 4D). This suggested that Path 4 represented macrophages that could become either M1 or M2. Interestingly, Path 4 showed higher *ATF3* and *ZNF384* TFs activity compared to other paths (Figure 4E). Paths 5 and 6 showed higher signal of unspliced reads for M1 and M2 markers, respectively, corroborating that these cells are likely terminally polarized M1 and M2. Our results indicated that there is cell to cell variability in response to microenvironmental stimuli and that our macrophage subpopulations are heterogeneous. In particular, we found a subpopulation that does not respond to stimulus and tends to maintain an M0 state, a very plastic subpopulation that can polarize towards both M1 and M2, and a subset that responds to microenvironmental stimuli and terminally polarizes towards either M1 or M2.

### Subpopulation of terminally polarized M2 macrophages express proliferation genes not observed in the M1 population

We sought to characterize the identity of our single-cell populations (Figure 4A) based upon expression of unique HL-60, M0, M1, and M2 markers obtained from our bulk RNA-seq analysis and from the literature. As expected, only Path 1 (HL-60-specific) presented high expression of *TOP2A*, which is a marker of aberrant cellular proliferation in cancer cells, and high levels of *TYMS*, which is known to be highly expressed in HL-60 cells (Pei et al., 2018; Ulger et al., 2003) (Figure S3B). M1 stimulation induced *PARP9* that has been shown to promote expression of pro-inflammatory genes (Iwata et al., 2016). Importantly, *P2RY14* gene that is involved in inflammatory signaling and induces cell cycle dormancy was present only in terminally polarized M1/Path 5, corroborating our findings that M1 cells do not continue cycling (Cho et al., 2014). M2 stimulation induced expression of genes *SLA* and *RASA3* which are involved in progression of cell cycle (Dulmovits et al., 2015) (Figure S3A). In addition, TF YY1 increases cell proliferation and is more active in M2 (Figure 4E). These results combined with ontology analysis of M2-specific genes identified in our bulk experiments (Figure 1E) suggest that M2 macrophages are capable of proliferating, whereas M1 macrophages might not be.

### M1 macrophages more strongly express polarization markers than M2

After quantification of spliced and unspliced single-cell reads, we observed that Path 5 (M1) expresses strong signals for both spliced and unspliced M1 markers, while Path 6 (M2) expresses weaker unspliced signals for M2 markers (Figure 4D). In addition, M1 showed more active TFs compared to M2 (Figure 4E). M1 stimulus led to increased activity of canonical M1 TFs STAT1, STAT2, IRF1, IRF7, NF_k_B1, and NF_k_B2. M2 stimulus promoted increased activity of TFs YY1, RUNX1/2 and MEF2A/C. These results suggest that M1 and M2 polarization are regulated by a distinct subset of TFs that is reflected at the single-cell level.

We explored orthogonal expression of M1 and M2 markers in individual single cells by calculating Odds-Ratio (OR) (see Methods) to further survey cell to cell variability in response to stimuli. For instance, we inquired whether a cell expressing a M1 marker X would have reduced probability (Negative OR, p value < 0.05) or increased probability (High/Positive OR, p value < 0.05) of expressing a M2 marker Y. We also investigated which pair of M1 and M2 markers are more likely to be expressed in the same cell (Figures 4F & S3C). We observed that M1 cells (Path 5) expressing M1-specific chemokines *CXCL10, CXCL9* and signaling molecule *TNFSF13B* are significantly less likely to express M2 markers (Figure 4F). Similarly, M1 cells that express M1-specific genes *GBP2, IFIT3*, and *FCGR1A* are less likely to express M2 markers (Figure 4F). When it comes to M2 cells (Path 6), the pair-wise expression correlations are not as strong (Figure S3C), as previously reported (Muñoz-Rojas et al., 2021). This suggests that terminally polarized M1 present a stronger phenotype and are less likely to express M2 markers.

### M2 repolarized towards M1 presents a unique transcriptome profile

We addressed whether a polarization switch from M1 to M2 or from M2 to M1 was possible to further evaluate macrophage plasticity and ability to repolarize. Remarkably, we were able to repolarize M2 cells towards an M1 subtype but were unable to repolarize M1 towards an M2 subtype, likely due to a stronger M1 phenotype and low likelihood of expressing M2 genes (Figure 4F). We applied fluorescence-activated cell sorting (FACS) to select terminally polarized M2, CD163 high M2 macrophages (SM2) in order to obtain a homogeneous M2 population (Hu et al., 2017). We further repolarized these cells with M1 stimuli LPS and IFN-γ (see Methods). We performed single-cell RNA-seq on M2 repolarized M1 (M2M1). We compared the transcriptome profiles of M2M1 cells with HL-60, M0 in addition to M1, M2 and SM2 (24 hours post-stimuli) and observed the clustering of subpopulations using UMAP (Figure 5A). We identified four distinct cell populations or paths (Figures 5A & S3D). As expected, HL-60 cells were enriched in a unique Path 1R and M0 cells were enriched in Path 2R (Figures 5B & S3E). Path 3R was mainly enriched in M1 and M2M1 cells and Path 4R was enriched in M2 and SM2. Next, we visualized the main markers identified in each population (Figure 5C). M2M1 showed a similar expression pattern as M1 macrophages, suggesting that the populations are transcriptionally very similar. Correspondingly, M2 and SM2 presented similar expression patterns. Unsupervised *in-silico* trajectory reconstruction by Monocle demonstrated that M0 bifurcated into two distinct trajectories that contained mainly M2M1 and M1 or mainly M2 and SM2 (Figure 5D). Based on the number of M2M1 cells that clustered with M1 (Figure S3E) we determined that approximately 90% of terminally polarized M2 cells repolarized towards an M1 subtype upon LPS and IFN-γ activation. M2M1 expressed M1-specific markers very similarly to M1 macrophages (Figures 5E & 5F). In addition, M2M1 showed high signal of unspliced M1 markers, suggesting that their future state is more M1-like (Figure 5F). Although M2M1 expressed M1-specific markers, they retained some unique transcriptional differences. M2M1 cells still expressed M2 genes associated with proliferation, such as *CDK1, MCM2*, and *MKI67* (Figure 5F). In our model, we observed that M2M1 macrophages acquired an M1-like transcriptome profile but likely maintained M2’s ability to proliferate.

**Figure 5.**
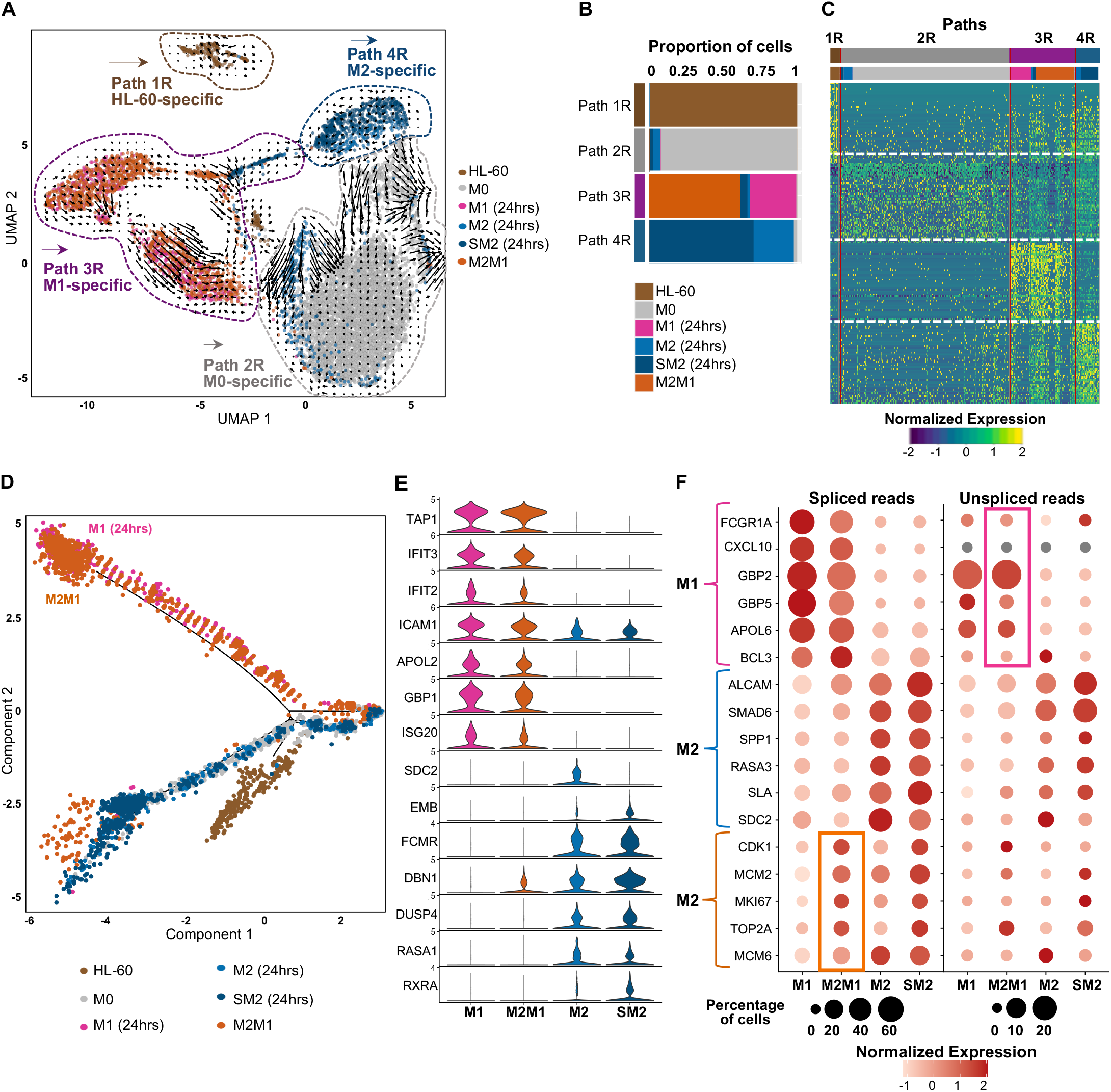
M2 repolarized towards M1 presents a unique transcriptome profile. A)UMAP embedding representation of single-cell RNA-seq repolarization time course annotated by time points and clusters of subpopulations (Paths). RNA velocity vectors were projected onto the UMAP and indicate future cellular trajectories. Brown, grey, pink, light blue, dark blue, and orange represent HL-60, M0, M1, M2, sorted M2 (SM2), and M2 repolarized towards M1 (M2M1) populations, respectively. B) Bar plot of the relative proportion of cell subtypes per cluster (Path). C) Heatmap of top differentially expressed genes per cluster. Each column represents a cluster of cells (Path), and each row represents a gene. Expression is row normalized. Column colors are as noted in B. D)Pseudotime trajectory of macrophage repolarization colored by time points E) Stacked violin plots of selected M1 and M2 markers. Plot shows markers’ expression per cluster. F)Bubble chart depicting spliced and unspliced reads for a given gene (row) per Path (column). M1- and M2-specific markers were highlighted in pink and blue, respectively. Orange brace highlights proliferation genes expressed in the M2, sorted M2 (SM2), and M2 repolarized towards M1 (M2M1) populations. Circle sizes represent the percentage of cells expressing the given gene. Color key represents normalized average expression.

### Single-cell GRN provides additional potential network connections

We built scATAC-seq libraries of subsets of M0, as well as 24-hour polarized M1 and M2 macrophages using the Bio-Rad ddSEQ platform to construct a single-cell GRN using scRNA-seq and scATAC-seq data sets (see Methods). We used the linked self-organizing maps (SOM) strategy (Jansen et al., 2019) to provide additional networking analysis from the single-cell data and compare the results to our bulk GRN. We built separate SOMs and performed metaclustering for each of the scRNA and scATAC datasets separately (Figure S4A). Metaclusters were linked by finding the closest gene within 1Mb for each genome region to create a multiclustering (see Methods). These linked metaclusters were each searched for motifs via FIMO (q-value < 0.05) and the motifs were filtered by linked metacluster enrichment (p value < 0.05). This provided a total of ∼500,000 unique network connections of which ∼36,000 were TF-TF interactions. We identified 135 connections between the 23 genes of our core network (Figure 6A). We recovered previously studied interactions including E2F4 regulation of *PU*.*1*/*SPI1* (Lachmann et al., 2010), PU.1 regulation of *CSF1* (Smith et al., 1996), as well as IRF7*’s* and MEF2A’s auto-regulatory loops (Ning et al., 2005; Ramachandran et al., 2008) (Table S1). The predicted auto-regulatory loop for *IRF1* was also an interesting finding considering the well-known feed-forward loop behavior seen in the IRF1-*STAT1* regulatory system (Michalska et al., 2018).

**Figure 6.**
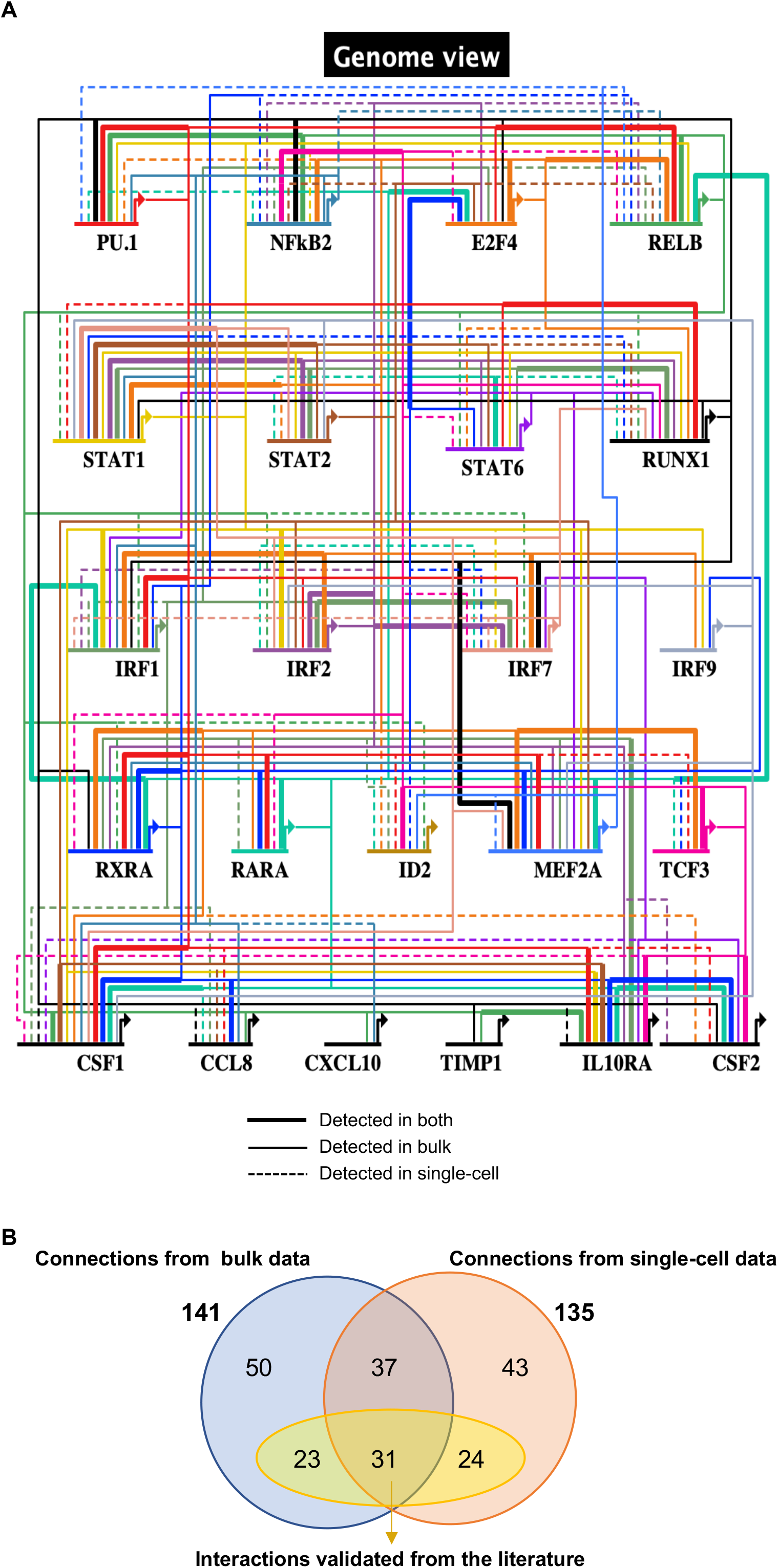
Comparison of single-cell-derived connections and bulk-derived connections. A) Gene regulatory networks containing 17 transcription factors and 6 signaling molecules. Genome view shows connections identified in the single-cell GRN, in the bulk GRN and connections identified by both methods. Solid bold lines indicate connections detected using both single-cell and bulk methods; solid lines indicate connections detected using the bulk method only; dashed lines indicate connections identified using the single-cell method only. Each color is specific to a transcription factor that regulates multiple targets. B)Venn diagram of total number of connections detected using bulk analysis (blue) and single-cell analysis (orange) as well as the overlaps between both methods. Yellow diagram highlights the connections confirmed by previous studies.

Footprinting analysis of bulk data identified a total of 141 connections, whereas SOM linking analysis of single-cell data identified 135 connections (Figure 6B). Comparison of the SOM linking-derived connections to those identified from the bulk analysis found that around half of these connections (68) overlapped. Overall, our single-cell-derived GRN identified 55 known interactions and 80 novel candidate interactions that regulate macrophage polarization, whereas our bulk-derived GRN identified 54 known interactions and 87 novel candidate interactions that regulate macrophage polarization. However, neither approach recovered all previously validated interactions from the literature, and the combined network is more complete than either strategy alone.

### Validation of GRN connections using siRNA in M2 polarization

We decided to confirm some of our M2 polarization GRN predictions using siRNA knockdown (KD) of four transcription factors of interest – IRF1, IRF7, IRF9, and ID2 (Figure 7A). We chose to perturb M2 since it has been less studied, and our predicted targets presented unique links that are M2-specific. For instance, *IRF1* is targeted by RARA only in M2; *IRF7* is targeted by PU.1, IRF1, and IRF2 only in M2 (Figures 3C & 6A). Therefore, we perturbed IRF1, IRF7, and IRF9 to explore their regulatory roles during M2 polarization. Moreover, we sought to explore which genes were affected by ID2^KD^ since our results showed that it might play an important role in M2 polarization.

**Figure 7.**
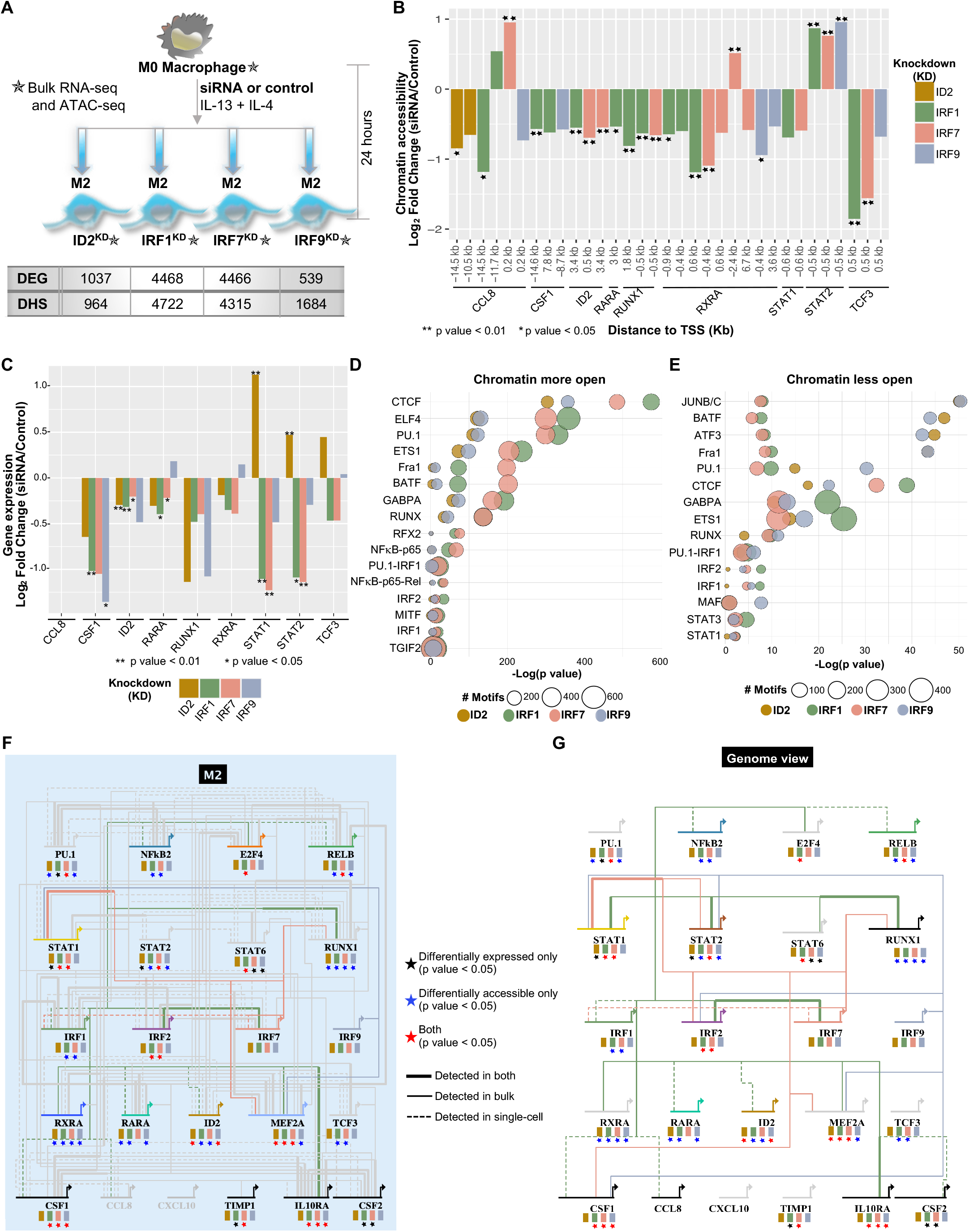
IRF1, IRF7, IRF9, and ID2 regulate distinct subsets of transcription factors during M2 polarization. A) Perturbation assays pipeline: M0 macrophages were treated with IRF1, IRF7, IRF9, ID2, or control siRNAs combined with M2-specific stimuli (IL-13 and IL-4). Cells were harvested 24 hours post-siRNA treatment. Table indicates the number of differentially expressed genes (DEG) and the number of differentially open regions of chromatin (DNase I hypersensitive sites - DHS) per knockdown. B) Chromatin accessibility fold change (log_2_) between ID2^KD^, IRF1^KD^, IRF7^KD^, IRF9^KD^, and control conditions (log_2_FC > 0.5) is shown for a partial list of regions associated with genes (15kb+/-) depicted in the gene regulatory networks of macrophage polarization. The distance from the chromatin element to the start of each gene is indicated. Differential accessibility is indicated ∗p value < 0.05, ∗∗p value < 0.01. C) Gene expression fold change (log_2_) between knockdowns and control is shown for a partial list of genes depicted in the gene regulatory networks of macrophage polarization. Differential expression significance was calculated using biological replicates (see Methods). ∗p value < 0.05, ∗∗p value < 0.01. D) Motif enrichment in regions that became more accessible upon ID2^KD^, IRF1^KD^, IRF7^KD^, and IRF9^KD^. Circle sizes represent the number of motifs for the given transcription factor (y axis). Each color represents the TF that was knocked down. E) Motif enrichment in regions that became less accessible upon ID2^KD^, IRF1^KD^, IRF7^KD^, and IRF9^KD^. Circle sizes represent the number of motifs for the given transcription factor (y axis). Each color represents the TF that was knocked down. F) Bulk M2-focused GRN (Figure 3C) centered on knockdown TFs. Links originating directly from IRF1, IRF7, or IRF9 are colored by green, pink, or blue, respectively, while all other links are in grey for simplicity. Target genes are colored if the single-cell analysis predicted a regulatory connection between the knockdown TF and the target gene. Any link present in both the bulk and the single-cell networks is in bold. The banners underneath each gene correspond to each knockdown (ID2 in gold, IRF1 in green, IRF7 in pink, IRF9 in blue). The stars represent the significance of the knockdown in both RNA-seq and ATAC-seq for that gene. Black stars indicate the gene is differentially expressed upon knockdown (p < 0.05), blue stars indicate differential chromatin accessibility determined by at least 1 region in 15kb+/-TSS (p value <0.05), and red stars indicate the gene is both differentially expressed and accessible. G) Single-cell-focused GRN (Figure 6A) centered on knockdown TFs. Target genes are colored if the single-cell analysis predicted a regulatory connection between the knockdown TF and the target gene. Differential expression and accessibility for each gene are noted as in F. Regulatory connection lines and colors are in the same style as in F.

We profiled bulk gene expression and chromatin accessibility after 24 hours of siRNA and M2 polarization stimuli (see Methods). We observed that chromatin became significantly less accessible in genomic regions associated with our KD targets *IRF1, IRF7, IRF9* and *ID2* (Figures 7B & S5A). Similarly, the expression of *IRF1, IRF7, IRF9*, and *ID2* was reduced compared to controls, confirming the successful knockdown (Figures 7C & S5B). ID2^KD^ downregulated 927 genes and upregulated 110 genes (Figure S5C). ID2 is known to repress *TCF3* and its expression increased in ID2^KD^ M2 (Figure 7C). Importantly, ID2^KD^ led to overexpression of M1-specific genes, such as *STAT1, STAT2*, and *NF*_*k*_*B2* suggesting that ID2 might repress important M1 markers during M2 polarization (Figures 7C & S5B). Differential motif enrichment calculated in regions whose chromatin accessibility changed upon knockdown showed that ID2^KD^ and IRF9^KD^ affected regions that contain similar motifs (Figures 7D & 7E). The chromatin regions that became less open upon ID2 reduction or IRF9 reduction were enriched in JUNB/C, ATF3, and BATF3 motifs (Figure 7E).

The number of chromatin regions that became more or less open upon IRF1^KD^ was very similar to the number for IRF7^KD^ (Figure S5D). In addition, chromatin regions that became more open upon IRF1^KD^ or IRF7^KD^ contained similar motifs, such as CTCF, ELF4, and PU.1 (Figure 7D). These results suggest that IRF1 and IRF7 might have similar regulatory roles during M2 macrophage polarization (Figures 7D & 7E). We found that several predicted IRF1 targets identified in our GRN showed significant changes in expression and/or chromatin accessibility upon *IRF1* knockdown. Such potentially-confirmed targets include *IRF2, IRF7, RUNX1*, and *IL10RA*, which were predicted to be IRF1 targets in M2 in both bulk and single-cell GRNs, as well as *RXRA* and *MEF2A* (detected in the bulk GRN), and *CSF1* and *ID2* (detected in the single-cell GRN) (Figures 7F & 7G). Similarly, IRF7 was predicted to target *RUNX1* (detected in the bulk GRN) and *IRF1* (detected in the single-cell GRN), which both showed changes in chromatin accessibility and/or gene expression upon IRF7 reduction. Some differences in expression and chromatin accessibility upon knockdown of our targets can also be explained by indirect TF regulation. *PU*.*1* expression significantly changes upon *IRF1* knockdown (Figure 7F). Although we did not identify *PU*.*1* as a direct IRF1 target, MEF2A is a predicted IRF1 target (detected in the bulk GRN) that seems to regulate *PU*.*1* expression (detected in the single-cell GRN) (Figures 6A & 7F). *MEF2A* expression significantly decreased in our IRF1^KD^ in M2, thus potentially explaining *PU*.*1* expression changes. Also, IRF7^KD^ led to changes in gene expression and chromatin accessibility in *IL10RA*, which is not a direct IRF7 target. However, STAT1 is a potential IRF7 target whose expression and chromatin accessibility change in IRF7^KD^ and therefore likely explains the expression changes of the STAT1 predicted target *IL10RA* (Figure 7F). In summary, our perturbations provided significant evidence to key connections identified in our bulk and single-cell GRNs by direct or indirect TF interactions.

## Discussion

Defining the gene regulatory networks that underlie cellular maturation in response to stimuli is a challenging task. A previous study used network modeling software on published gene expression data to identify macrophage polarization regulators (Palma et al., 2018). Although it highlighted important players in the M1/M2 activation, it did not identify the main TF-target interactions that underlie the process. Our bulk footprinting-derived GRN revealed distinct and subtype-specific TF-target connections between 23 polarization genes. M1 had 40 unique connections, and M2 had 17 unique connections; 63 connections were shared between M1 and M2, suggesting a large concordance between the M1 and M2 polarization pathways. In addition, some polarization markers seem to play distinct roles in both M1 and M2 GRNs. We captured the known activation of *IRF1* by STAT6 in the M1 GRN, but not in M2 (Miller et al., 2019). *STAT6* is a well-known marker of M2, while *IRF1* is a pro-inflammatory marker of M1. Our identified connections implied that STAT6 may still have a regulatory role in driving a pro-inflammatory response in M1 that is absent in M2. Around 32% (23 of 73) of our bulk-specific interactions have been confirmed in previous studies, and 68% are novel candidate interactions.

As regulatory genomics is rapidly adopting single-cell methods, we sought to verify whether single-cell techniques would reproduce or improve our bulk RNA-seq and ATAC-seq derived networks. Our single-cell GRN identified 135 connections, of which 68 overlapped our bulk GRN. It identified 67 connections not found using bulk data, whereas bulk analysis identified 73 connections not found in the single-cell GRN. Similar to our bulk GRN, around 36% (24 of 67) of our single-cell-specific interactions have been confirmed by the literature and 64% are novel candidate interactions. Around 46% (31 of 68) of the links predicted by both methods have been confirmed in previous studies, which is a higher percentage compared to each method separately.

Using our bulk data sets, we detected rapid changes in gene expression as early as 3 hours post polarization. Large chromatin accessibility changes during polarization were detected later in the time course, similar to results seen during macrophage differentiation (Ramirez et al., 2017). *FCGR1A*, whose expression is increased in immuno-inflammatory syndromes (Minar et al., 2014), reacted to M1, but not M2 stimuli as early as 3 hours of polarization. Therefore, *FCGR1A* is a possible target to modulate M1 polarization for therapeutics of inflammatory disorders (Thepen et al., 2009). In turn, *CSF2* reacted to M2 stimuli but not M1 as early as 3 hours. *CSF2* is a potential candidate target to alter M2 polarization through modulation of autophagy and could possibly be a target for cancer therapeutics (Chen et al., 2014). Overall, our results provide early, intermediary and late polarization markers that could facilitate the manipulation of macrophages to desired polarized states and could therefore reveal novel therapeutic targets for cancers and immune diseases.

ID2 is an important regulator of macrophage development *in vivo* and *in vitro*. ID2^KD^ mice lacked specialized skin macrophages, called Langerhans cells (Hacker et al., 2003). However, ID2 has never been explored as a macrophage polarization marker. Given its increased expression and chromatin accessibility during our M2 polarization time course, we suggest that it can be a novel marker of M2 polarization. Additionally, we found that ID2^KD^ led to overexpression of M1 markers suggesting it may induce the M2 lineage by suppressing M1 markers.

M0 macrophages’ ability to proliferate has been well explored (Daems & De Bakker, 1982), but it is less clear whether M1 or M2 can proliferate. Skin resident macrophages are able to proliferate in order to maintain the tissue population throughout life (Hacker et al., 2003). However, during inflammatory conditions, macrophages were unable to proliferate and were instead replaced by circulating monocytes (Dieu-Nosjean et al., 2000). In our model, M1 expressed genes linked to cell cycle dormancy suggesting that M1 did not enter the cell cycle, similar to what has been seen in inflammatory contexts. On the other hand, our M2 cells expressed proliferation genes not seen in M1 and thus seemed to proliferate. A previous study has shown that macrophage proliferation is a hallmark of T Helper 2 signaling and is linked to IL-4, which corroborates our findings (Jenkins et al., 2011). These results suggested that macrophage proliferation is directly linked to microenvironmental stimuli. Thus, naive M0 and M2 stimulated by IL-4 are able to proliferate, whereas M1 stimulated by inflammatory signals are not. Notably, despite acquiring an M1-like transcriptome, our M1 cells that repolarized from M2 maintained expression of proliferative genes. Therefore, M1 cells that were once M2 may inherit proliferative capabilities.

Both bulk and single-cell functional genomics methods used in our studies have advantages and limitations that will shape the resulting networks. The bulk footprinting method we used is known to show high efficiency, but it relies on protein occupancy, which may not detect TFs with short-lived binding. Hence, important polarization TFs such as NF_k_B, which are very dynamic, do not always leave footprints in bulk chromatin (Sung et al., 2016). In contrast, single-cell assays face technical challenges, including resolution, sparsity and number of genes detected. This last limitation is present in our analysis due to 3’ tagmentation methods that produce dropouts. For example, we were unable to detect *PU*.*1* reliably in our scRNA-seq experiments, which severely restricted our ability to detect its connections in the single-cell GRN. Therefore, neither method alone was able to predict all known interactions, and more complete results were obtained by a combination of both bulk and single-cell GRNs.

In summary, our findings provide a clearer understanding of the heterogeneity of the cells that undergo M1 and M2 polarization and M2 to M1 repolarization. We identified transient polarization markers and found that ID2 is a novel M2 TF. We constructed *de novo* GRNs containing both known and novel interactions that underlie M1 and M2 polarization. It will be interesting to investigate how much of this polarization network is conserved across heterogeneous tissue-resident macrophages in health and disease, especially in the context of microglial polarization in AD. In conclusion, we believe that despite the revolution and the great potential of single-cell biology, bulk results are still needed for building more comprehensive and accurate GRNs where practical, i.e., where we can isolate relatively pure cell types for bulk assays. We expect that insights gained from this work will provide experimental and computational guidelines for building GRNs of cellular maturation in response to microenvironmental stimuli.

## Acknowledgments

This work was supported in part by grants from the National Institutes of Health UM1HG009443 and R01AG060148 to A.M and by a fellowship from CNPq/LASPAU.

## Author Contributions

A.M. and K.C. conceived the study. K.C. performed macrophage differentiation, polarization, and repolarization, RNA-seq and ATAC-seq libraries preparation and sequencing, scRNA-seq libraries preparation, as well as analysis of RNA-seq, ATAC-seq, and scRNA-seq data; E.R performed scATAC-seq libraries preparation and bulk footprinting analysis; C.J. performed SOM-linking analysis; K.W. performed odds-ratio analysis; A.D. maintained HL-60 cell culture; C.M. sequenced the scRNA-seq and scATAC-seq libraries. All authors contributed to writing of the manuscript.

## Methods

### CONTACT FOR REAGENT AND RESOURCE SHARING

#### Lead Contact

Further information and requests for reagents may be directed to, and will be fulfilled by the Lead Contact Ali Mortazavi.

#### Materials Availability

This study did not generate new materials.

### EXPERIMENTAL MODEL AND SUBJECT DETAILS

HL-60 cells (ATCC-CC240) were grown in ATCC-recommended media: 20% FBS (Omega Scientific) and 1% penicillin/streptomycin antibiotics (Life Technologies) in Dulbecco’s Modification of Eagle’s Medium (Corning). They were incubated at 37°C with 5% CO_2_. All cell lines were maintained in horizontally oriented 25mL or 75mL Falcon Tissue Culture Treated Flasks (Thermo Fisher Scientific) at a density of 1 × 10^6^ cells/mL in a total of 10mL or 20mL, respectively. Cells were consistently passaged once over 2 to 3-day periods up to differentiation.

### METHOD DETAILS

#### Macrophage differentiation and polarization

We performed PMA-induced differentiation of HL-60 cells into M0 macrophages (Murao et al., 1983) in order to obtain M1 and M2 macrophage subtypes for characterization. Approximately 5 × 10^6^ cells at a density of 1 × 10^6^ cells/mL were plated in 60mm cell culture dishes with 10 µM of PMA (Thermo Fisher Scientific). Media was changed every 48 hours and 10µM of PMA was added at every media change. After 120 hours of PMA stimulus, M0 macrophages were polarized by either applying 100ng/mL of IFN-γ and LPS to obtain M1 or 10ng/mL of IL-4 and IL-13 to obtain M2 polarized subtypes (Huang et al., 2018; Wang et al., 2007). Differentiation and polarization were confirmed by fluorescence immunostaining for cell type specific markers (Rőszer, 2015; Yu et al., 2009; Holness & Simmons, 1993) and by observation of markedly distinct cellular morphologies, as M1 cells are characterized by a large cell body with a varied number of pseudopodia and M2 cells are characterized by an elongated shape (McWhorter et al., 2013). Macrophages were collected at 0, 3, 6, 12 and 24 hours after addition of polarization stimuli, along with undifferentiated HL-60 for bulk and single-cell RNA-seq (Figure 1A). We generated a total of 62 bulk data sets, approximately 50,000 single-cells using scRNA-seq, and approximately 40,000 single-nuclei using scATAC-seq.

#### M2 repolarization towards M1

M0 macrophages were polarized towards M2 for 24 hours according to aforementioned protocol. M2 macrophages were harvested and resuspended in PBS at a concentration of 1 × 10^4^ cells/mL. Non-specific antigens were blocked by 5% BSA in PBS buffer. Cells were then fluorescently labeled using CD163-FITC mouse conjugated monoclonal antibody (Thermo Fisher, MA5-17719). Cells that presented high CD163 levels were sorted using an Aria 2 Flow Cytometer (BD Biosciences) in order to obtain a homogeneous M2 population (Hu et al., 2017). M2 macrophages were repolarized towards M1 by applying 100ng/mL of IFN-γ and LPS during 24 hours. Populations of M2 repolarized towards M1 (M2M1) were harvested in a single-cell suspension and loaded into the ddSEQ single-cell isolator (Bio-Rad). Single-cell libraries were prepared using the SureCell WTA 3’ Library Prep Kit for the ddSEQ System (Illumina). The quality of the libraries was assessed using the Agilent 2100 Bioanalyzer. Single-cell libraries were sequenced using the NextSeq 500 (Illumina). Cell viability (>90%) was confirmed prior to single-cell library preparation.

#### Bulk RNA-seq and ATAC-seq experiments

Bulk sequencing experiments were conducted in triplicates per time point (Figure 1A) per assay. We collected 2 million cells for RNA-seq and 50,000 cells for ATAC-seq. RNA-seq and corresponding ATAC-seq biological replicates were collected from the same dish. Cell viability (>90%) was monitored prior to cell collection. Macrophages were detached from plates by incubating with 3.5mL of Trypsin-EDTA 0.25% (Life Technologies) at 37°C for 4 minutes. Trypsin was neutralized by adding 14mL of complete media per dish, then the cells were pelleted in a centrifuge at 1,500 RPM for 5 minutes. The cells were washed with PBS and spun down again to remove all traces of Trypsin and media before library preparation. HL-60 cells were collected for RNA-seq by pelleting suspended cells and washing them with PBS. We relied on our previous paper for undifferentiated HL-60 ATAC-seq data (Ramirez et al., 2017). RNA-seq and ATAC-seq libraries were built following the Smart-seq2 protocol (Picelli et al., 2014) and the Omni-ATAC protocol (Corces et al., 2017), respectively. The Omni-ATAC-seq libraries went through a gel size selection step to enrich for DNA fragments ranging from 150 to 500bp. The quality of all libraries was assessed using the Agilent 2100 Bioanalyzer. Bulk libraries were sequenced using the NextSeq 500 (Illumina) obtaining around 10 million reads per sample for RNA-seq and around 20 million reads per sample for ATAC-seq. Two replicates per time point were used in downstream analyses.

#### Single-cell RNA-seq experiment

Single-cell libraries were prepared from 12,000 cells per time point (Figure 1A) using the SureCell WTA 3’ Library Prep Kit for the ddSEQ System (Illumina). Cells with viability >90% were collected by detaching cells from plates using Trypsin-EDTA 0.25%, as previously described in the “Bulk RNA-seq and ATAC-seq experiments” section. The detached cells were washed in cold PBS + 0.1% BSA and resuspended to reach 2,500 cells/µL. The cells were filtered through a 40 µM strainer and verified to be in single-cell suspensions under the microscope. Cells suspended in a reverse transcription reaction buffer were loaded into the Bio-Rad ddSEQ microfluidic device along with barcoded beads. Single cells were encapsulated in an oil-water emulsion as nanodroplets and their RNA was reverse transcribed, during which unique molecular identifiers (UMIs) and cell barcodes were added to the cDNA. The first strand cDNA was recovered from the droplets through emulsion breakage and bead cleanup, then the second strand of cDNA was synthesized. The cDNA was cleaned, tagmented, amplified, and cleaned again to produce the final libraries. The concentration and size distribution of the libraries were assessed using the Agilent 2100 Bioanalyzer. Single-cell libraries were sequenced using the NextSeq 500 (Illumina) at 3.1 pM loading concentration using a custom primer.

#### Single-cell ATAC-seq experiment

The Omni-ATAC version of the Bio-Rad ddSEQ SureCell ATAC-seq (Illumina) workflow was followed, based off of the bulk ATAC-seq protocol of the same name (Lareau et al., 2019; Corces et al., 2017). Single-cell suspensions of M0, as well as 24-hour polarized M1 and M2 samples were prepared as previously described in the “Single-Cell RNA-seq experiment” section. Cells were resuspended to approximated 1×10^6^/mL in PBS + 0.1% BSA, filtered through a 40µM strainer, and verified to be single-cell suspensions under the microscope. Samples of 300,000 cells per biological replicate of M0, M1, and M2 were lysed with ATAC-Lysis buffer containing freshly added digitonin, then the nuclei were washed with ATAC-Tween buffer. Staining with Trypan Blue (Bio-Rad) confirmed cells were lysed with viability <10% and nuclei were verified to be single-nucleus suspensions under the microscope. 60,000 nuclei were used in each Tn5 tagmentation reaction (incubation at 37°C for 30 min) per sample. After tagmentation, nuclei suspended in an amplification reaction were loaded into the Bio-Rad ddSEQ microfluidic device along with barcoded beads. Single nuclei were encapsulated in nanodroplets and the fragments generated by the Tn5 transposase were barcoded by UMIs and cell barcodes and amplified. The fragments were recovered from the droplets through emulsion breakage, bead cleanup, a second amplification reaction, and a final bead cleanup before quality control using the Agilent 2100 Bioanalyzer. Libraries were assessed by concentration and size distribution before being loaded onto the NextSeq 500 (Illumina) and sequenced at 1.5 pM loading concentration using a custom primer.

#### IRF1, IRF7, IRF9 knockdown in M2

M0 cells were differentiated from HL-60 then polarized to M2 for 24 hours as previously described in the “Macrophage differentiation and polarization” section. At the same time that polarization reagents were added, siRNAs targeting IRF1, IRF7, IRF9, ID2, positive control (GAPD), and negative (non-targeting) control (Dharmacon) were transfected in 3 biological replicates of M2 per target knockdown. DharmaFECT 4 Transfection Reagent was added at 1:500 final concentration and siRNAs were added at 0.025µM final concentration in 4mL media in 60mm cell culture dishes. The perturbed cells were collected for both bulk RNA-seq and ATAC-seq 24 hours later.

### QUANTIFICATION AND STATISTICAL ANALYSIS

#### Bulk RNA-seq preprocessing

Bulk RNA-seq reads were mapped to the hg38 reference genome with gene annotations from Gencode release 29 using STAR (version 2.5.1b) and gene level expression was quantified using RSEM (version 1.2.25) (Dobin et al., 2013; Li & Dewey, 2011). Expression represented by counts per gene was normalized utilizing the “weighted” Trimmed Mean of M-values (TMM) approach using edgeR (version 3.28.1) and saved as a matrix of TMM normalized counts per million (CPM) (Robinson & Oshlack, 2010). Counts were also converted to transcripts per million (TPM).

#### Bulk RNA-seq analysis

EdgeR (version 3.28.1) also identified genes differentially expressed between selected time points using a false discovery rate (FDR) of 1% and an alpha of 0.05. TMM normalized CPM were then log_2_ normalized and used as input for maSigPro (version 1.58.0) to identify gene expression changes over time allowing for k-means clustering of genes that present similar patterns of expression during the time course of polarization.

#### Bulk ATAC-seq preprocessing

ATAC-seq reads were mapped to the hg38 genome, annotated by Gencode release 29, using Bowtie2 (version 2.2.7) (Langmead & Salzberg, 2012). Reads mapped to the mitochondrial chromosome were discarded from downstream analysis. The resulting bam file was sorted using Picard toolkit (version 2.18.4) and PCR duplicates were removed (https://broadinstitute.github.io/picard/). Using a custom script, the remaining reads aligning to the positive strand were shifted by +4bps and reads aligning to the negative strand were shifted by −5bps to account for the 9bp duplication caused by Tn5 transposition (Berg et al., 1983). We used HOMER (version 4.7) to create a tag directory for each sample using the sorted and shifted bam file (Heinz et al., 2010). To determine the final set of open regions: peaks were called for both “narrow” regions of 150bp and “broad” regions of 500bp using HOMER (version 4.7) and the resulting bed files were merged across replicates for each sample. Biologically reproducible regions across replicates were identified by irreproducible discovery rate analysis (IDR), then ENCODE “blacklist” regions were removed in order to generate the final set of open regions (Amemiya et al., 2019; Li et al., 2011). We used HOMER (version 4.7) to estimate read coverage across the final set of open regions using the tag directories generated for each sample, and to annotate each region.

#### Bulk ATAC-seq analysis

ATAC-seq read counts were corrected for TMM and library size (represented as CPM) using edgeR (version 3.28.1) to normalize the data. EdgeR was also used to identify differentially open chromatin regions within gene promoters at each of the differentiation time points. TMM normalized counts were then log_2_ transformed and maSigPro (version 1.58.0) was used to cluster regions based on changes in accessibility during the polarization time course (Nueda et al., 2014). All clusters were converted into bed files of genomic regions and HOMER (version 4.7) findMotifsGenome.pl was used to find which known motifs were most enriched in each cluster.

#### Bulk data integration and gene regulatory network construction

We applied Pearson’s χ2 test from the R Stats Package (version 3.6.2) to determine p values of enrichment between ATAC-seq clusters and RNA-seq clusters. A final list of TFs was manually curated by combining TFs found in RNA clusters, linked RNA-ATAC cluster pairs, results from scRNA-seq analysis, motif enrichment in ATAC clusters, and differentially expressed TFs across the time course. ATAC-seq data across all the M1 or M2 replicates from 3 hours to 24 hours were pooled in order to achieve >200 million reads to mine the open regions for transcription factor footprints. We used the Wellington algorithm from the pyDNase library (version 0.2.4) and the HINT-ATAC tool to identify chromatin footprints with an FDR of 1%, with footprint sizes of 8 to 31bp with a 1bp step size (Li et al., 2019; Piper et al., 2013). We then used FIMO (version 4.12.0) to identify footprint regions with enriched HOCOMOCO (version 11) transcription factor motifs, using a p value cutoff of 0.0001 and hg38 background (Kulakovskiy et al., 2013). We used bedtools intersect to determine the target genes for each footprint using the HOMER-annotated bulk ATAC-seq regions. We finally used footprints <50kb away from the TSSs of the target genes to build bulk GRNs with our TFs and signaling molecules of interest.

#### Single-Cell RNA-seq preprocessing

Single-cell RNA-seq reads were demultiplexed using ddSeeker, which extracts cell barcodes and UMIs and produces a demultiplexed bam file (Romagnoli et al., 2018). From the bam file, we used a custom script to extract one fastq file per cell. Each cell was mapped to the hg38 reference genome with gene annotations from Gencode release 29 using STAR (version 2.5.1b). Spliced and unspliced reads were calculated using velocyto (version 0.17.16) and expression was also quantified using RSEM (version 1.2.25) (La Manno et al., 2018). Low quality cells with less than 500 UMI counts, more than 20% mitochondrial reads, and less than 150 genes detected were removed from further analysis. We also removed cells with more than 4,000 genes detected to avoid possible doublets.

#### Single-cell RNA-seq analysis

Each cell that passed the initial quality control filters was used as input for Velocyto (La Manno et al., 2018). Velocyto (version 0.17.16) only considers uniquely mapped reads that align to both exonic and intronic regions and removes reads mapped to repeat masked regions. The new UMI count matrices were exported from a loom file format to a Seurat object (V3.1.5) (Satija et al., 2015). Downstream normalization, differential expression and Leiden clustering were implemented using Seurat (Traag et al, 2019). RNA velocities calculated by Velocyto were then overlaid onto the Seurat UMAP dimensionality reduction. To further investigate cell trajectories we used the R package Monocle (version 2.8.0) to order cells within a pseudotime utilizing an unsupervised clustering method (Qiu et al., 2017). We then applied Single-Cell Regulatory Network Inference And Clustering (SCENIC, version 1.1.2-2) to estimate the transcription factors that regulate gene expression on cell populations identified using Seurat (Aibar et al., 2017). We finally applied the Odds-Ratio (OR) analysis to investigate orthogonal expression of M1 and M2 markers in individual cells. The odds-ratio was calculated from the spliced counts. Genes with greater than 2 counts were considered “expressed”. The odds-ratio was calculated as the product of the number of cells expressing both genes (YY) and the number of cells expressing neither gene (NN) divided by the product of the number of cells expressing only one gene (YN) and the number of cells expressing only the other gene (NY). OR=(YYxNN)/(YNxNY). Plots use log_2_ of the odds-ratio with any infinite values set to the positive or negative absolute maximum value. All odds-ratio values with a p value > 0.05 were set to 0.

#### Single-cell ATAC-seq preprocessing

The average number of nuclei collected and average number of unique fragments per nucleus between technical replicates of M0 were 2,480 and 4,104, respectively. For two sets of biological replicates with two technical replicates each (4 libraries) of M1, we recovered 3,860 nuclei and 8,102 unique fragments per cell on average. For two sets of biological replicates with two technical replicates of M2, we recovered 4,404 nuclei and 8,839 unique fragments on average. Raw sequencing data were primarily processed using the Bio-Rad ATAC-seq Analysis Toolkit (version 1.0.0) in order to generate mapped reads. We installed the Bio-Rad Docker containers on our lab’s server and followed the steps to perform fastq QC, debarcoding of fastq files, alignment to hg38 using BWA with blacklist regions removed, and alignment QC using Picard Tools (Li & Durbin, 2009). Next, the Bio-Rad Toolkit used Jaccard indexing to determine bead duplicates and remove cell-free droplets, then BAP to “deconvolute” or merge bead duplicates. The mapped reads for UMI-passing cells and the peaks called by MACS2 (Zhang et al., 2008) were used in a custom python script (found here: https://github.com/fairliereese/lab_pipelines/tree/master/sc_atac_pipeline) to generate peaks-by-cells counts matrices for each sample for use in SOM analysis.

#### Single-cell SOM analysis

To analyze the single-cell data, we trained a self-organizing map on the read counts for each of the 210,312 fragments for each cell/nucleus. For this, we used our SOMatic tool (Jansen & Ramirez et al., 2019) with a 40×60 map for 5 epochs and 100 trials. This process resulted in 114 DNA metaclusters. Similarly, we built a 40×60 map on the scRNA data with the same options and received 52 RNA metaclusters. Then, these 2 sets of metaclusters were linked, generating 5928 (114 × 52) linked metaclusters. The regions in these linked metaclusters were sent through the same network analysis pipeline as our previous work (above) using the HOCOMOCOv11 motif database (p value < 0.05, q-value < 0.05). The pipeline can be found here: https://github.com/csjansen/SOMatic-Network-Analysis. This generated a total of 8,904,925 potential network connections or 833,114 TF-TF interactions.

### DATA AND SOFTWARE AVAILABILITY

The accession number for the sequencing data reported in this paper is GEO: GSE164498.

