## Supplemental Material for "Uncovering the Gene Regulatory Networks Underlying Macrophage Polarization Through Comparative Analysis of Bulk and Single-Cell Data"

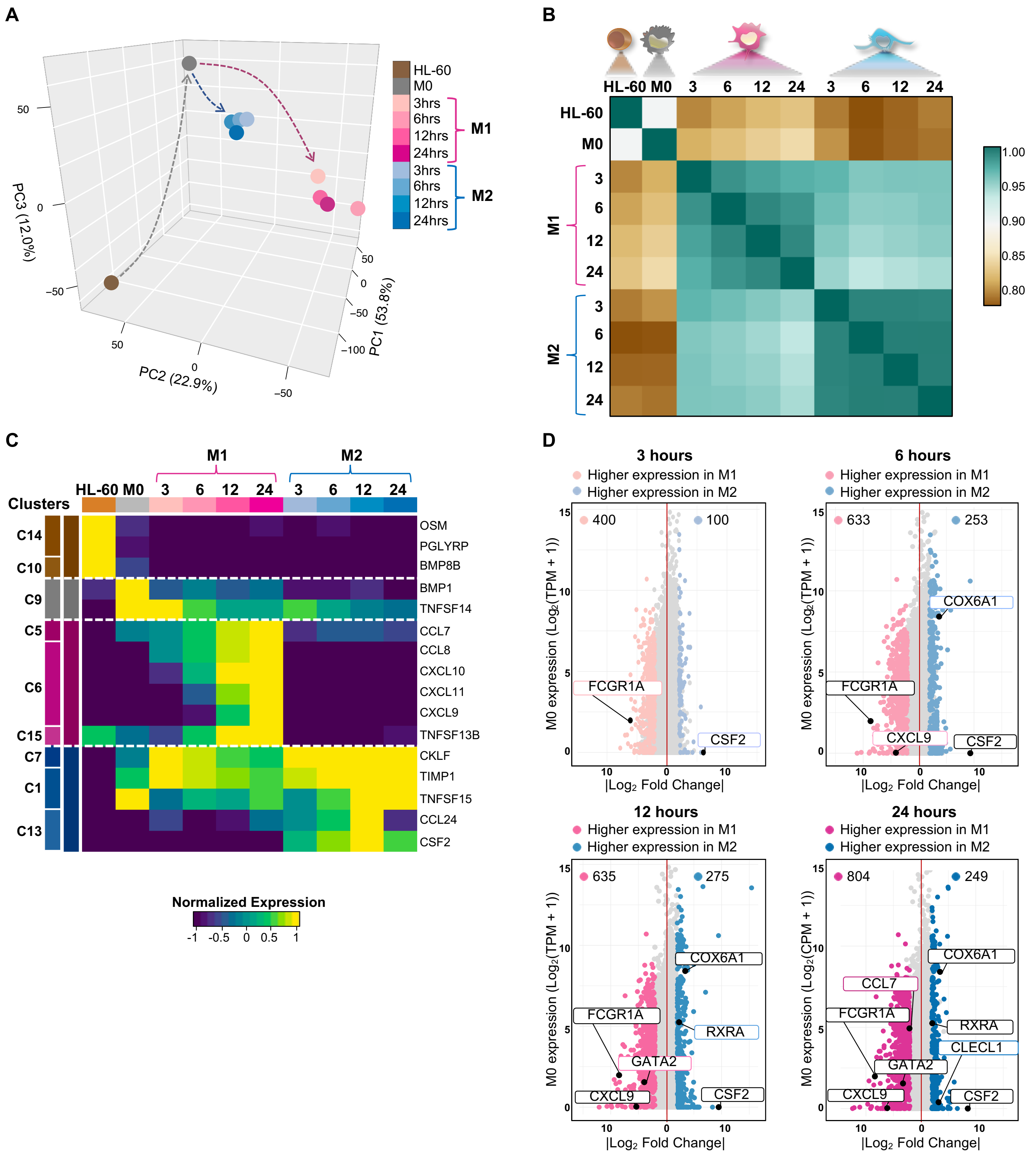


**Figure S1. Related to Figure 1. Characterizing temporal changes in gene expression during macrophage polarization**

A) Principal component analysis of the RNA-seq time course of macrophage polarization. Time points were connected to illustrate subtype-specific trajectories. Cell types are depicted in distinct colored points.

B) Pearson correlation analysis of the RNA-seq time course of macrophage polarization.

C) Heatmap of signaling molecules included in HL-60-, M0-, M1- or M2-specific clusters identified by MaSigPro. Each column represents the average expression for a time point and each row represents a signaling molecule. RNA-seq data is row-mean normalized.

D) Genes differentially expressed between M1 and M2 time points ($log_2FC > 1$, FDR < 0.05) compared to normalized M0. M1- and M2-specific genes FCGR1A, CSF2, CXCL9, COX6A1, GATA2, RXRA, CCL7, and CLECL1 are indicated.

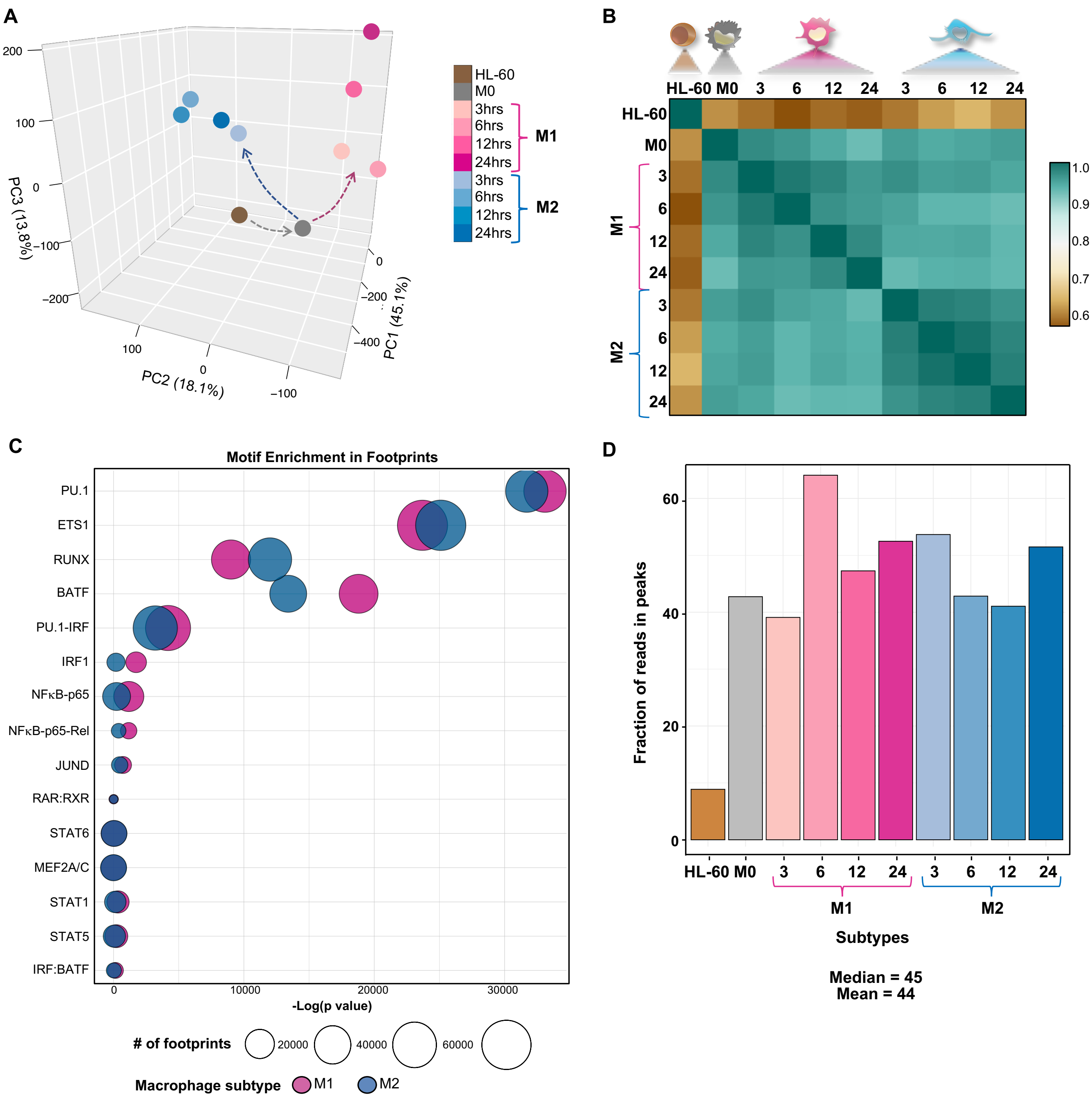


**Figure S2. Related to Figure 2. Quality assessment of bulk ATAC-seq data**
A) Principal component analysis of the ATAC-seq time course of macrophage polarization. Time points were connected to illustrate subtype-specific trajectories. Cell types are depicted in distinctly colored points.
B) Pearson correlation analysis of the ATAC-seq time course of macrophage polarization**.**
C) Differential motif enrichment between chromatin accessibility footprints in M1 and M2 subtypes compared to each other. Circle sizes represent the number of footprintings for the given transcription factor.
D) Distribution of ATAC-seq fraction of reads in peaks (FRiP). Mean = 44, median = 45.

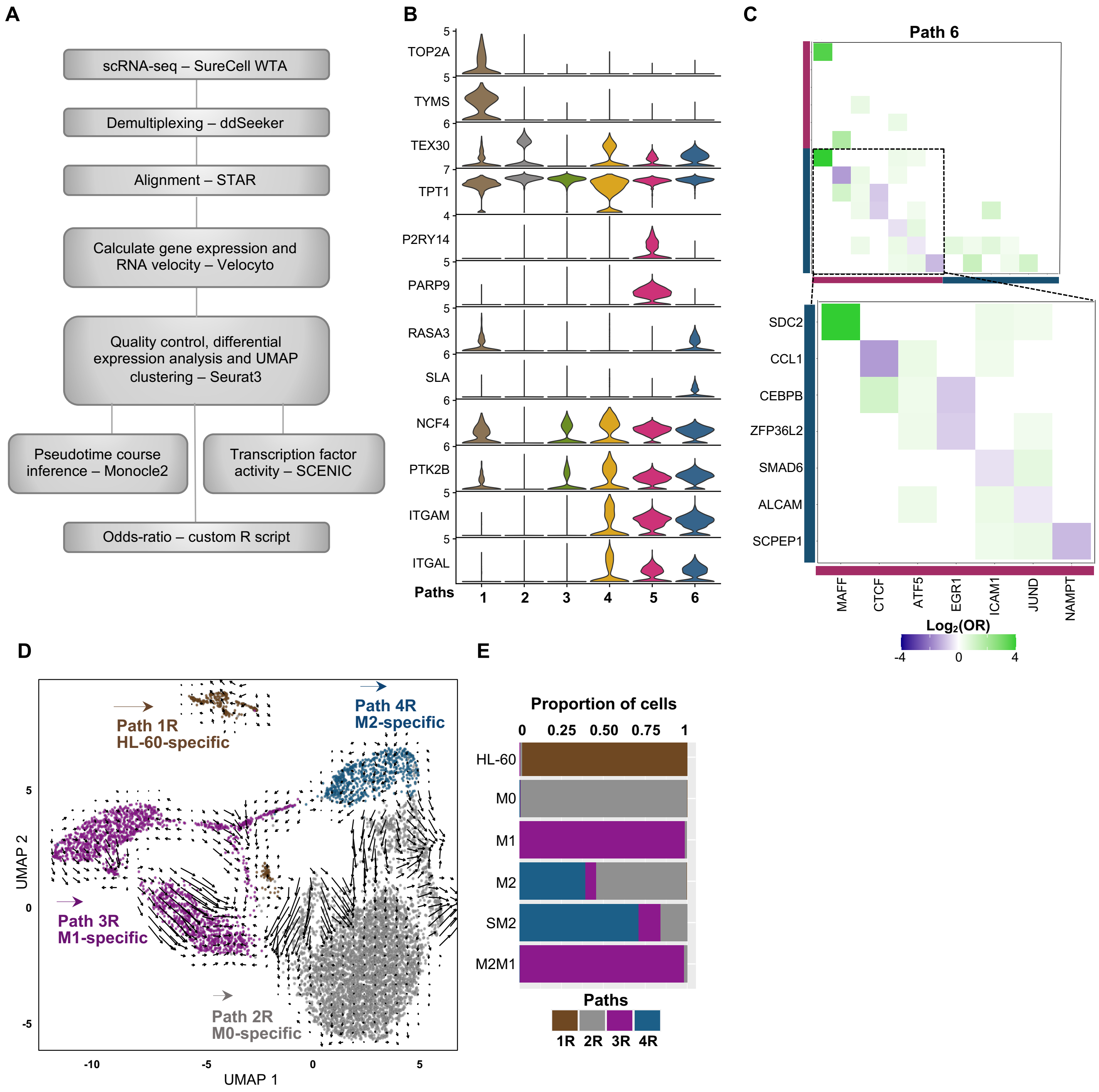


**Figure S3. Related to Figures 4 and 5. Single-cell RNA-seq of macrophage polarization and repolarization**

A) Single-cell RNA-seq data analysis pipeline.

B) Stacked violin plots of selected markers' expression per cluster (Path).

C) Pairwise odds ratios (OR) for M1 genes (pink) and M2 genes (blue) detected in Path 6. Odds ratios that present a p value > 0.05 (calculated by Fisher's exact test) are set to 0. Data shown as $\log_2$(odds ratio).

D) UMAP embedding representation of the single-cell RNA sequencing repolarization time course annotated by clusters of subpopulations. RNA velocity vectors were projected onto the UMAP and indicate future cellular trajectories.

E) Bar plot of the relative proportion of cell subtypes per cluster.

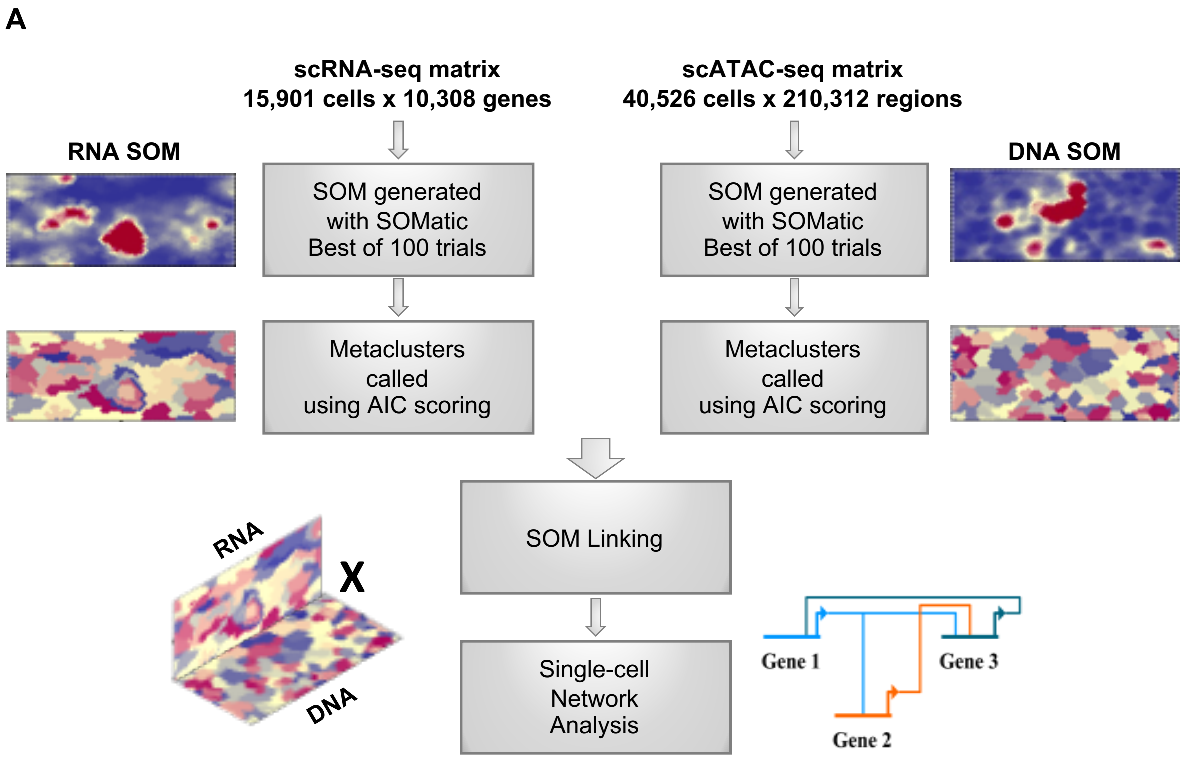


**Figure S4. Related to Figure 6. Single-cell RNA-seq and ATAC-seq data integration using SOM linking**
A) SOM linking data analysis pipeline used to build the single-cell gene regulatory networks.

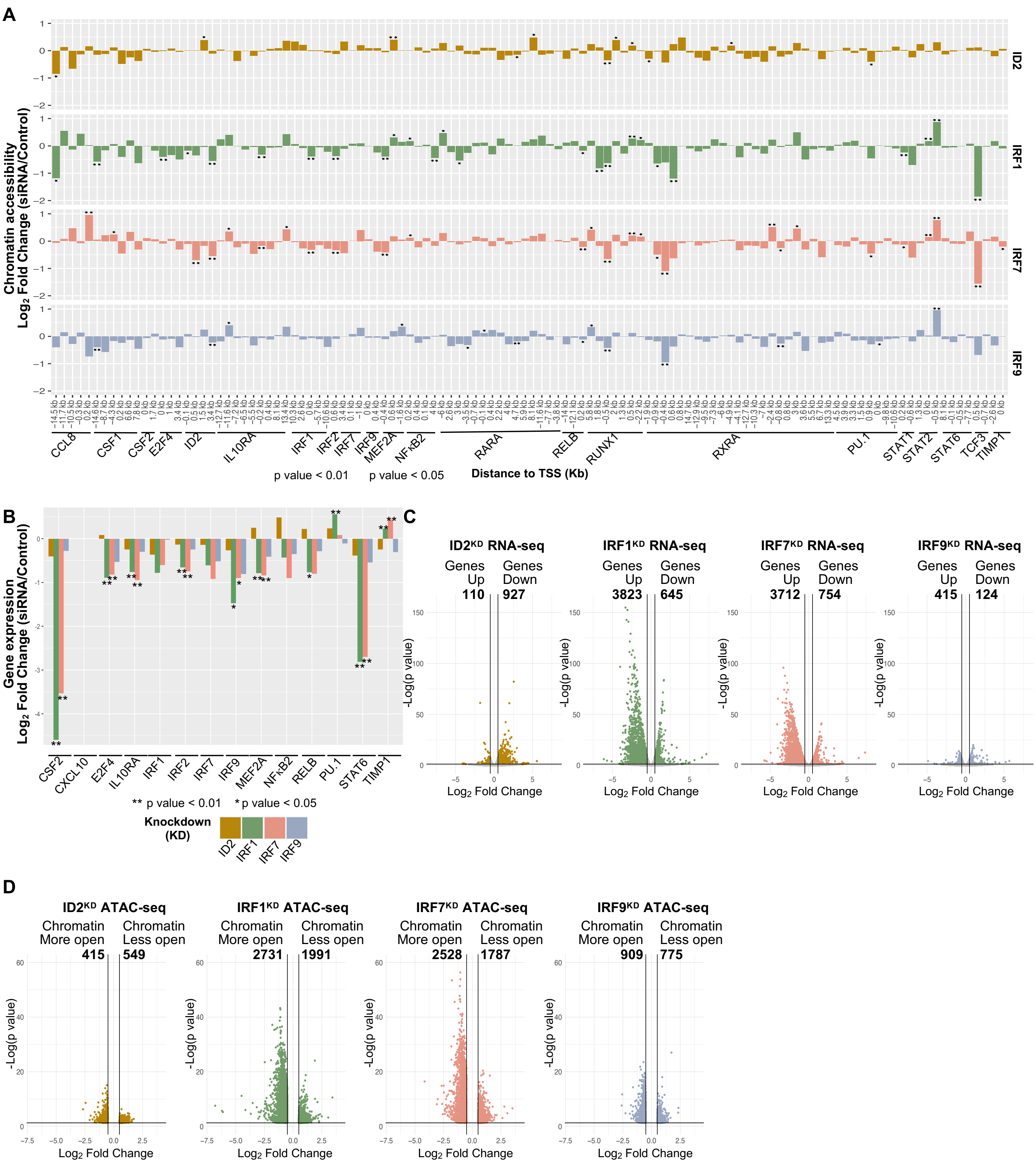


**Figure S5. Related to Figure 7. Differential gene expression and chromatin accessibility upon $IRF1^{KD}$, $IRF7^{KD}$, $IRF9^{KD}$, and $ID2^{KD}$**

A) Chromatin accessibility fold change ($\log_2$) between $ID2^{KD}$, $IRF1^{KD}$, $IRF7^{KD}$, $IRF9^{KD}$, and control conditions is shown for regions associated with genes (15kb+/-) depicted in the gene regulatory networks of macrophage polarization. The distance from the chromatin element to the start of each gene is indicated. Differential accessibility is indicated $*p$ value < 0.05, $**p$ value < 0.01.

B) Gene expression fold change ($\log_2$) between knockdowns and control is shown for a partial list of genes depicted in the gene regulatory networks of macrophage polarization. Differential expression significance was calculated using biological replicates (see Methods). $*p$ value < 0.05, $**p$ value < 0.01.

C) Volcano plot highlighting genes upregulated or downregulated in $ID2^{KD}$, $IRF1^{KD}$, $IRF7^{KD}$, $IRF9^{KD}$. Differentially expressed genes are colored in gold, green, orange, and blue, respectively. p value < 0.05, $|log_2FC| > 0.5$.
D) Differential chromatin accessibility between $ID2^{KD}$, $IRF1^{KD}$, $IRF7^{KD}$, $IRF9^{KD}$, and control conditions. Differentially open regions are colored in gold, green, orange, and blue, respectively. p value < 0.05, $|log_2FC| > 0.5$.

### Table S1: Related to Figure 6. Regulatory interactions that were previously studied

| Citations | Interactions | Identified in our data |
|---|---|---|
| Bonizzi, G., Bebien, M., Otero, D. C., Johnson-Vroom, K. E., Cao, Y., Vu, D., … Karin, M. (2004). Activation of IKKα target genes depends on recognition of specific κB binding sites by RelB:p52 dimers. EMBO Journal, 23(21), 4202–4210. https://doi.org/10.1038/sj.emboj.760039 | RELB auto-regulatory loop | Both |
| Bren, G. D., Solan, N. J., Miyoshi, H., Pennington, K. N., Pobst, L. J., & Paya, C. V. (2001). Transcription of the RelB gene is regulated by NF-κB. Oncogene, 20(53), 7722–7733. https://doi.org/10.1038/sj.onc.1204868 | RELB->IRF7 | Single-cell |
| De Silva, N. S., Anderson, M. M., Carette, A., Silva, K., Heise, N., Bhagat, G., & Klein, U. (2016). Transcription factors of the alternative NF-kB pathway are required for germinal center B-cell development. Proceedings of the National Academy of Sciences of the United States of America, 113(32), 9063–9068. https://doi.org/10.1073/pnas.1602728113; Lovas, A., Radke, D., Albrecht, D., Buket, Z. B., Möller, U., Habenicht, A. J. R., & Weih, F. (2008). Differential RelA- and RelB-dependent gene transcription in LTβR-stimulated mouse embryonic fibroblasts. BMC Genomics, 9, 606. https://doi.org/10.1186/1471-2164-9-606 | RELB->STAT6 | Single-cell |
| Dong, X., Craig, T., Xing, N., Bachman, L. A., Paya, C. V, Weih, F., … Griffin, M. D. (2003). Direct transcriptional regulation of RelB by 1α,25-dihydroxyvitamin D3 and its analogs: physiologic and therapeutic implications for dendritic cell function. Biochemistry, (33). | RXRA->NFkB2 | Single-cell |
| El Zein, R. M., Soria, A. H., Golib Dzib, J. F., Rickard, A. J., Fernandes-Rosa, F. L., Samson-Couterie, B., … Boulkroun, S. (2019). Retinoic acid receptor α as a novel contributor to adrenal cortex structure and function through interactions with Wnt and Vegfa signalling. Scientific Reports, 9(1), 14677. https://doi.org/10.1038/s41598-019-50988-2 | RARA->TCF3 | Single-cell |
| Fijneman, R. J. A., Anderson, R. A., Richards, E., Liu, J., Tijssen, M., Meijer, G. A., … Cormier, R. T. (2012). Runx1 is a tumor suppressor gene in the mouse gastrointestinal tract. Cancer Science, 103(3), 593–599. https://doi.org/10.1111/j.1349-7006.2011.02189.x | RUNX1->CCL8 | Single-cell |
| Ho, J., Pelzel, C., Begitt, A., Mee, M., Elsheikha, H. M., Scott, D. J., & Vinkemeier, U. (2016). STAT2 Is a Pervasive Cytokine Regulator due to Its Inhibition of STAT1 in Multiple Signaling Pathways. PLOS Biology, 14(10), e2000117. https://doi.org/10.1371/journal.pbio.2000117 | STAT2->STAT1 | Both |
| Hohaus, S., Petrovick, M. S., Voso, M. T., Sun, Z., Zhang, D. E., & Tenen, D. G. (1995). PU. 1 (Spi-1) and C/EBP alpha regulate expression of the granulocyte-macrophage colony-stimulating factor receptor alpha gene. Molecular and cellular biology, 15(10), 5830-5845. DOI: 10.1128/MCB.15.10.5830 | PU.1->CSF2 | Single-cell |
| I, A., WY, Y., A, F., JJ, C., EA, W., & MR, H. (2014). p75NTR is highly expressed in vestibular schwannomas and promotes cell survival by activating nuclear transcription factor κB. Glia, 62(10). https://doi.org/10.1002/GLIA.22709 | TCF3->NFkB2 | Both |
| Ji, M., Li, H., Suh, H. C., Klarmann, K. D., Yokota, Y., & Keller, J. R. (2008). Id2 intrinsically regulates lymphoid and erythroid development via interaction with different target proteins. Blood, The Journal of the American Society of Hematology, 112(4), 1068-1077. https://doi.org/10.1182/blood-2008-01-133504 | PU.1->ID2 | Single-cell |
| Koh, C.P., Wang, C.Q., Ng, C.E.L., Ito, Y., Araki, M., Tergaonkar, V., Huang, G., and Osato, M. (2013). RUNX1 meets MLL: epigenetic regulation of hematopoiesis by two leukemia genes. Leukemia 27, 1793–1802. | RUNX1->MEF2A | Both |
| Kwon, G., & Kang, K. (2020). Transcriptional regulation of IL10 gene in human macrophages. | PU.1->IL10RA | Both |
| Lachmann, A et al. (2010) ChEA: transcription factor regulation inferred from integrating genome-wide ChIP-X experiments. Bioinformatics. 26:2438-44. - Harmonizome - D R, GW G, NF F, Z W, CD M, et al. (2016) The harmonizome: a collection of processed datasets gathered to serve and mine knowledge about genes and proteins. Database(Oxford) Jul 3;2016 | E2F4->NFkB2 | Both |
| Laslo, P., Spooner, C.J., Warmflash, A., Lancki, D.W., Lee, H.J., Sciammas, R., Gantner, B.N., Dinner, A.R., and Singh, H. (2006). Multilineage Transcriptional Priming and Determination of Alternate Hematopoietic Cell Fates. Cell 126, 755–766; Kubosaki, A., Tomaru, Y., Tagami, M., Arner, E., Miura, H., Suzuki, T., Suzuki, M., Suzuki, H., and Hayashizaki, Y. (2009). Genome-wide investigation of in vivo EGR-1 binding sites in monocytic differentiation. Genome Biology 10, R41 | PU.1 auto-regulatory loop | Both |

| | | |
|---|---|---|
| Lee et al., (2011) Wide-ranging functions of E2F4 in transcriptional activation and repression revealed by genome-wide analysis. Nucleic Acids Research, Volume 39, Issue 9, https://doi.org/10.1093/nar/gkq1313 | E2F4->IRF1<br>E2F4->IRF2<br>E2F4->IRF7<br>E2F4->MEF2A<br>E2F4->STAT1<br>E2F3->TCF3 | Both |
| Lie-A-Ling, M., Marinopoulou, E., Li, Y., Patel, R., Stefanska, M., Bonifer, C., … Lacaud, G. (2014). RUNX1 positively regulates a cell adhesion and migration program in murine hemogenic endothelium prior to blood emergence. Blood, 124(11), 11–20. https://doi.org/10.1182/blood-2014-04-572958 | RUNX1->PU.1 | Both |
| Lovas, A., Radke, D., Albrecht, D., Buket, Z. B., Möller, U., Habenicht, A. J. R., & Weih, F. (2008). Differential RelA- and RelB-dependent gene transcription in LTβR-stimulated mouse embryonic fibroblasts. BMC Genomics, 9, 606. https://doi.org/10.1186/1471-2164-9-606 | RELB->ID2 | Single-cell |
| Mathelier, A et al. (2014) JASPAR 2014: an extensively expanded and updated open-access database of transcription factor binding profiles. Nucleic Acids Res. 42:D142-7. - Harmonizome - D R, GW G, NF F, Z W, CD M, et al. (2016) The harmonizome: a collection of processed datasets gathered to serve and mine knowledge about genes and proteins. Database(Oxford) Jul 3;2016 | IRF2->STAT1 | Both |
| Michalska et al., (2018) A Positive Feedback Amplifier Circuit That Regulates Interferon (IFN)-Stimulated Gene Expression and Controls Type I and Type II IFN Responses. Frontiers in Immunology, https://doi.org/10.3389/fimmu.2018.01135 | IRF1->STAT1 | Both |
| Minucci, S., Leid, M., Toyama, R., Saint-Jeannet, J. P., Peterson, V. J., Horn, V., ... & Ozato, K. (1997). Retinoid X receptor (RXR) within the RXR-retinoic acid receptor heterodimer binds its ligand and enhances retinoid-dependent gene expression. Molecular and Cellular Biology, 17(2), 644-655. DOI: 10.1128/MCB.17.2.644 | RXRA->RARA | Both |
| Nair, P. M., Starkey, M. R., Haw, T. J., Ruscher, R., Liu, G., Maradana, M. R., … Hansbro, P. M. (2018, March 1). RelB-deficient dendritic cells promote the development of spontaneous allergic airway inflammation. American Journal of Respiratory Cell and Molecular Biology, Vol. 58, pp. 352–365. https://doi.org/10.1165/rcmb.2017-0242OC | RELB->PU.1 | Both |
| Nakagawa, M., Shimabe, M., Watanabe-Okochi, N., Arai, S., Yoshimi, A., Shinohara, A., … Kurokawa, M. (2011). AML1/RUNX1 functions as a cytoplasmic attenuator of NF-κB signaling in the repression of myeloid tumors. Blood, 118(25), 6626–6637. https://doi.org/10.1182/blood-2010-12-326710 | RUNX1->IRF7 | Both |
| Nakanishi, M., Tomaru, Y., Miura, H., Hayashizaki, Y., & Suzuki, M. (2008). Identification of transcriptional regulatory cascades in retinoic acid-induced growth arrest of HepG2 cells. Nucleic Acids Research, 36(10), 3443–3454. https://doi.org/10.1093/nar/gkn066 | RARA->IRF1 | Both |
| | RARA->STAT6 | Both |
| | RXRA->RUNX1 | Single-cell |
| | RXRA->TCF3 | Single-cell |
| Navarro-Montero, O., Ayllon, V., Lamolda, M., López-Onieva, L., Montes, R., Bueno, C., … Real, P. J. (2017). RUNX1c Regulates Hematopoietic Differentiation of Human Pluripotent Stem Cells Possibly in Cooperation with Proinfla | RUNX1->CSF1 | Single-cell |
| Nigten, J., Nikoloski, G., De Witte, T., Van der Reijden, B. A., & Jansen, J. H. (2004). Id1 and Id2 Are Retinoic Acid Responsive Genes and Induce a G0/G1 Arrest in Acute Promyelocytic Leukemia Cells. https://doi.org/10.1182/blood.V104.11.2029.2029 | RARA->ID2 | Single-cell |
| Ning, S., Huye, L. E., & Pagano, J. S. (2005). Regulation of the transcriptional activity of the IRF7 promoter by a pathway independent of interferon signaling. Journal of Biological Chemistry, 280(13), 12262-12270. doi: 10.1074/jbc.M404260200 | IRF7 auto-regulatory loop | Single-cell |
| Orlikova, B., Schnekenburger, M., Zloh, M., Golais, F., Diederich, M., & Tasdemir, D. (2012). Natural chalcones as dual inhibitors of HDACs and NF-κB. Oncology Reports, 28(3), 797. https://doi.org/10.3892/OR.2012.1870 | STAT1->NFkB2 | Single-cell |
| Pongubala, J. M. R., & Atchison, M. L. (1997). PU.1 can participate in an active enhancer complex without its transcriptional activation domain. Proceedings of the National Academy of Sciences of the United States of America, 94(1), 127. https://doi.org/10.1073/PNAS.94.1.127 | PU.1->TCF3 | Single-cell |
| Ramachandran, B., Yu, G., Li, S., Zhu, B., & Gulick, T. (2008). Myocyte enhancer factor 2A is transcriptionally autoregulated. Journal of Biological Chemistry, 283(16), 10318-10329. doi: 10.1074/jbc.M707623200 | MEF2A auto-regulatory loop | Single-cell |
| Wislet, S., Vandervelden, G., & Rogister, B. (2018). From Neural Crest Development to Cancer and Vice Versa: How p75NTR and (Pro)neurotrophins Could Act on Cell Migration and Invasion?. Frontiers in molecular neuroscience, 11, 244. https://doi.org/10.3389/fnmol.2018.00244 | TCF3 auto-regulatory loop | Both |
| Saeed, S., Logie, C., Stunnenberg, H. G., & Martens, J. H. A. (2011). Genome-wide functions of PML-RARα in acute promyelocytic leukaemia. British Journal of Cancer, 104(4), 554–558. https://doi.org/10.1038/sj.bjc.6606095 | RARA->RUNX1<br>RARA->PU.1 | Single-cell |
| Satoh, J. I., Asahina, N., Kitano, S., & Kino, Y. (2014). A comprehensive profile of ChIP-Seq-based PU. 1/Spi1 target genes in microglia. Gene regulation and systems biology, 8, GRSB-S19711. https://doi.org/10.4137/GRSB.S19711 | PU.1->CSF1 | Both |
| | PU.1->STAT1 | Single-cell |

| | | |
|---|---|---|
| Sin, W.-X., Yeong, J. P.-S., Lim, T. J. F., Su, I.-H., Connolly, J. E., & Chin, K.-C. (2020). IRF-7 Mediates Type I IFN Responses in Endotoxin-Challenged Mice. Frontiers in Immunology, 11, 640. https://doi.org/10.3389/fimmu.2020.00640 | STAT1->IRF7 | Single-cell |
| Tallam, A., Perumal, T. M., Antony, P. M., Jäger, C., Fritz, J. V., Vallar, L., ... & Michelucci, A. (2016). Gene regulatory network inference of immunoresponsive gene 1 (IRG1) identifies interferon regulatory factor 1 (IRF1) as its transcriptional regulator in mammalian macrophages. PLoS One, 11(2), e0149050. https://doi.org/10.1371/journal.pone.0149050 | IRF1->RUNX1 | Both |
| The FANTOM Consortium., Suzuki, H., Forrest, A. et al. The transcriptional network that controls growth arrest and differentiation in a human myeloid leukemia cell line. Nat Genet 41, 553–562 (2009). https://doi.org/10.1038/ng.375 | IRF1->IRF2 | Both |
| | IRF2 auto-regulatory loop | Both |
| | PU.1->IRF1 | Both |
| | PU.1->RUNX1 | Both |
| Thomsen, I., Kunowska, N., Souza, R. de, Moody, A.-M., Crawford, G., Wang, Y.-F., … Sabbattini, P. (2018). RUNX1 controls the dynamics of cell cycle entry of naïve resting B cells by regulating expression of cell cycle and immunomodulatory genes in response to BCR stimulation. E-Conversion - Proposal for a Cluster of Excellence.; Navarro-Montero, O., Ayllon, V., Lamolda, M., López-Onieva, L., Montes, R., Bueno, C., … Real, P. J. (2017). RUNX1c Regulates Hematopoietic Differentiation of Human Pluripotent Stem Cells Possibly in Cooperation with Proinfla | RUNX1->IL10RA | Single-cell |
| Tomaru, Y., Simon, C., Forrest, A. R. R., Miura, H., Kubosaki, A., Hayashizaki, Y., & Suzuki, M. (2009). Regulatory interdependence of myeloid transcription factors revealed by matrix RNAi analysis. Genome Biology, 10(11), R121. https://doi.org/10.1186/gb-2009-10-11-r121 | RXRA->RELB | Single-cell |
| Wan, Y. J. Y., Wang, L., & Wu, T. C. J. (1994). The expression of retinoid X receptor genes is regulated by all-trans-and 9-cis-retinoic acid in F9 teratocarcinoma cells. Experimental Cell Research, 210(1), 56–61. https://doi.org/10.1006/excr.1994.1009; Nakanishi, M., Tomaru, Y., Miura, H., Hayashizaki, Y., & Suzuki, M. (2008). Identification of transcriptional regulatory cascades in retinoic acid-induced growth arrest of HepG2 cells. Nucleic Acids Research, 36(10), 3443–3454. https://doi.org/10.1093/nar/gkn066 | RARA->RXRA | Both |
| Waters, M. R., Gupta, A. S., Mockenhaupt, K., Brown, L. S. N., Biswas, D. D., & Kordula, T. (2019). RelB acts as a molecular switch driving chronic inflammation in glioblastoma multiforme. Oncogenesis, 8(6). https://doi.org/10.1038/s41389-019-0146-y | RELB->CSF1 | Both |
| | RELB->IRF2 | Single-cell |
| Wontakal, S. N., Guo, X., Will, B., Shi, M., Raha, D., Mahajan, M. C., ... & Skoultchi, A. I. (2011). A large gene network in immature erythroid cells is controlled by the myeloid and B cell transcriptional regulator PU. 1. PLoS Genet, 7(6), e1001392. ttps://doi.org/10.1371/journal.pgen.1001392 | E2F4->PU.1 | Single-cell |
| Zenke, K., Muroi, M., & Tanamoto, K. I. (2018). IRF1 supports DNA binding of STAT1 by promoting its phosphorylation. Immunology and Cell Biology, 96(10), 1095-1103. https://doi.org/10.1111/imcb.12185 | STAT1->IRF1 | Both |

### Table S2: Statistics for bulk and single-cell experiments

| | | Bulk RNA-seq | | Bulk ATAC-seq | | | scRNA-seq | | scATAC-seq | |
|---|---|---|---|---|---|---|---|---|---|---|
| Timepoint | Replicate | Mapped reads | Genes > 1TPM | Mapped reads | Homer total peaks | IDR-passing peaks | Cells > 500UMI | Cells >150 genes & <20% MT reads | Cells that pased QC | Avg. TSS Enrichment Score |
| HL-60 | Rep1 | 9.0M | 13,187 | 21.5M | 531,523 | 49,081 | 4,123 | 628 | – | – |
| | Rep2 | 8.4M | 13,124 | 14.1M | 737,361 | | | | | |
| Mac_0hrs | Rep1 | 13.9M | 14,436 | 15.8M | 514,659 | 131,631 | 4,480 | 4,145 | 4,960 | 42 |
| | Rep2 | 12.7M | 14,374 | 8M | 584,661 | | | | | |
| M1_3hrs | Rep1 | 13.5M | 10,369 | 29.3M | 584,661 | 168,480 | 635 | 415 | – | – |
| | Rep2 | 17.2M | 9,250 | 25.1M | 494,114 | | | | | |
| M1_6hrs | Rep1 | 13.5M | 10,473 | 19.7M | 431,602 | 162,497 | 3,453 | 475 | – | – |
| | Rep2 | 13.1M | 9,369 | 13.3M | 391,476 | | | | | |
| M1_12hrs | Rep1 | 11.7M | 9,744 | 25.3M | 482,322 | 170,366 | 571 | 490 | – | – |
| | Rep2 | 16.2M | 9,820 | 56.1M | 459,273 | | | | | |
| M1_24hrs | Rep1 | 13.2M | 9,385 | 19M | 508,889 | 164,340 | 597 | 524 | 8,425 | 40 |
| | Rep2 | 12.9M | 11,136 | 11.7M | 423,695 | | | | | |
| M2_3hrs | Rep1 | 8.0M | 8,228 | 15.3M | 451,124 | 157,267 | 474 | 435 | – | – |
| | Rep2 | 12.0M | 8,109 | 15.9M | 460,571 | | | | | |
| M2_6hrs | Rep1 | 10.0M | 7,722 | 16.1M | 501,319 | 148,226 | 507 | 389 | – | – |
| | Rep2 | 13.7M | 7,510 | 16.3M | 505,241 | | | | | |
| M2_12hrs | Rep1 | 9.5M | 7,413 | 23.1M | 462,814 | 153,577 | 877 | 285 | – | – |
| | Rep2 | 10.7M | 7,585 | 29.4M | 549,522 | | | | | |
| M2_24hrs | Rep1 | 9.0M | 7,788 | 20.3M | 506,421 | 187,605 | 494 | 418 | 8,809 | 37 |
| | Rep2 | 10.0M | 8,123 | 20.9M | 504,361 | | | | | |
| M2_24hrs_Sorted | – | – | | – | – | – | 829 | 630 | – | – |
| M2M1_Repolarized | – | – | | – | – | – | 1,323 | 1,159 | 329 | 16 |
| M2_Neg_Ctrl | Rep1 | 15.5M | 11,293 | 23.9M | 459,529 | 172,866 | – | – | – | – |
| | Rep2 | 18.3M | 12,022 | 24.4M | 459,889 | | | | | |
| M2_Pos_Ctrl | Rep1 | 13.1M | 12,373 | 25.9M | 479,147 | 112,537 | – | – | – | – |
| | Rep2 | 15.0M | 11,674 | 23.8M | 485,730 | | | | | |
| M2_IRF1_KD | Rep1 | 13.3M | 10,643 | 23.3M | 471,072 | 142,157 | – | – | – | – |
| | Rep2 | 15.5M | 10,669 | 27.3M | 514,258 | | | | | |
| M2_IRF7_KD | Rep1 | 13.7M | 10,534 | 27.9M | 508,087 | 166,015 | – | – | – | – |
| | Rep2 | 13.7M | 10,770 | 23.2M | 490,966 | | | | | |
| M2_IRF9_KD | Rep1 | 18.4M | 11,125 | 25.1M | 484,455 | 184,304 | – | – | – | – |
| | Rep2 | 15.8M | 11,545 | 25.3M | 477,066 | | | | | |
| M2_ID2_KD | Rep1 | 20.9M | 11,760 | 22.6M | 473,521 | 181,900 | – | – | – | – |
| | Rep2 | 10.8M | 12,016 | 23.2M | 465,532 | | | | | |
